# Genetic architecture of male courtship behavior differences in the parasitoid wasp genus *Nasonia* (Hymenoptera; Pteromalidae)

**DOI:** 10.1101/831735

**Authors:** J Gadau, C. Pietsch, S. Gerritsma, S. Ferber, L. van de Zande, J. van den Assem, L.W. Beukeboom

**Affiliations:** Institute for Evolution and Biodiversity, WWU Münster, Hüfferstraße 1, 48149 Münster; KWS SAAT AG, Grimsehlstrasse 31, 37555 Einbeck, Germany; Evolutionary Genetics, Centre for Ecological and Evolutionary Studies, University of Groningen, P.O. Box 11103, NL-9700 CC Groningen, The Netherlands

**Keywords:** Speciation, quantitative genetics, behavioral genetics, Quantitative Trait Locus, epistasis, prezygotic isolation, haplodiploidy

## Abstract

Very little is known about the genetic basis of behavioral variation in courtship behavior, which can contribute to speciation by prezygotic isolation of closely related species. Here, we analyze the genetic basis and architecture of species differences in the male courtship behavior of two closely related parasitoid wasps *Nasonia vitripennis* and *N. longicornis.* Both species occur microsympatrically in parts of their ranges and have been found in the same host pupae. Despite strong postzygotic isolation mechanisms between these two *Nasonia* species, viable hybrid females can be produced in the laboratory if both species are cured of their *Wolbachia* endosymbionts. We used haploid F_2_ hybrid males derived from virgin F_1_ hybrid females of two independent mapping populations to study the genetic architecture of five quantitative and two qualitative components of their courtship behavior. A total of 14 independent Quantitative Trait Loci (QTL) were found in the first mapping population (320 males), which explained 4-25% of the observed phenotypic variance. Ten of these QTL were confirmed by a second independent mapping population (112 males) and no additional ones were found. A genome-wide scan for two-loci interactions revealed many unique but mostly additive interactions explaining an additional proportion of the observed phenotypic variance. Courtship QTL were found on all five chromosomes and four loci were associated with more than one QTL, indicating either possible pleiotropic effects of individual QTL or individual loci contributing to multiple courtship components. Our results indicate that these two evolutionary young species have rapidly evolved multiple significant phenotypic differences in their courtship behavior that have a polygenic and highly interactive genetic architecture. Based on the location of the QTL and the published *Nasonia* genome sequence we were able to identify a series of candidate genes for further study.

## Introduction

Even closely related species can differ in conspicuous morphological, physiological or behavioral traits. Some of these differences can also promote speciation by blocking gene flow, i.e. acting either as post- or prezygotic isolation mechanisms. Several “speciation” genes involved in postzygotic isolation have been published over the last decade^1^ but very little is known about the genetic basis of prezygotic isolation^1,2^. Species differences in courtship behavior are important mechanisms of prezygotic isolation and there is evidence that behavioral traits leading to prezygotic isolation can evolve relatively rapidly if sister taxa occur sympatrically^3–6^. The speed with which this prezygotic isolation evolves depends on many factors. One crucial factor is the number and effect of genes (major, pleiotropic, epistatic) underlying the species differences in courtship behavior. Arbuthnott^2^ recently reviewed the literature on the genetic architecture of insect premating isolation and found that most reported courtship traits appear to be regulated by few loci with large effects. However, whether this result is evolutionarily relevant or is simply due to technical limitations of Quantitative Trait Locus (QTL) mapping (e.g. Beavis effect^7–9^) has recently been debated^10^. To resolve this requires tests of the effect of sample size on the probability to detect QTL with small effects and of overestimation of QTL effects in small mapping populations. We used two independent mapping populations that differed significantly in size to empirically test the effect of sample size on detecting QTL.

We investigated the genetic architecture of species differences in male courtship components between two closely related parasitoid wasp species, *Nasonia vitripennis* and *N. longicornis*^11–13^. *Nasonia* are small (2-3 mm) gregarious parasitoid wasps that parasitize fly pupae. The genus *Nasonia* consists of the cosmopolitan species *N. vitripennis* and three North-American species; *N. longicornis, N. giraulti* and *N. oneida*^14,15^. *N. vitripennis* occurs in some areas of its distribution sympatrically with the other three species. All species pairs, with the exception of the *N. oneida* - *N. giraulti* species pair, are reproductively isolated by cytoplasmic *Wolbachia* infections that result in nucleo-cytoplasmic incompatibilities^15,16^. Once cured of *Wolbachia*, *Nasonia* species are able to interbreed despite various degrees of postzygotic isolation mechanisms^17–20^

The *Nasonia* courtship fits the general pattern of courtship behavior within the Hymenopteran subfamily Pteromalinae^21^. Courtship behavior of *Nasonia* males is not learned and is composed of stereotyped motor patterns, which differ quantitatively and qualitatively between species^11,21^. This suggests that these components are genetically “hardwired” or based on allelic variation, which makes them suitable for QTL analysis. Courting males take up an invariable courtship position on top of the female, with the front legs placed on the female’s head^21^. A conspicuous feature of the courtship performance is head nodding: repeated bouts of nods that are separated by short pauses^11^. The first head nod in a bout coincides with mouthpart extrusions. Interspecific mate preference may also be asymmetric, i.e. females of one species refuse heterospecific males, whereas females of the other species accept conspecific as well as heterospecific males^12,13^.

QTL analysis enables us to estimate, in an unbiased manner (in contrast to a candidate gene approach), the number and effect of genes contributing to a given trait, as well as the interactions between these genes and hence is the method of choice to study the genetic basis of phenotypic differences between species. QTL analyses have been employed in a vast number of studies to investigate the genetic basis of species differences in morphological, behavioral and life-history traits and have substantially contributed to our understanding of the genetic architecture of species differences^22–32^. For example, interspecific crosses often produce individuals that have phenotypes not observed in either parental species (= transgressive phenotypes)^33^. There is some experimental evidence that alleles with opposing effects within each parental line provide the genetic basis for such transgressive segregation^34–35^ which can be evaluated using QTL studies. Another explanation that has been put forward to explain transgressive phenotypes is epistasis, i.e. the non-additive effect of a particular allele on the phenotype depending on the genetic background^33^. We also observed transgressive phenotypes for all of our traits and therefore analyzed the role of epistasis and alleles with opposing effects for the occurrence of transgressive phenotypes in our mapping population. The occurrence of epistasis in natural populations and hybrids is widely established, contributing for example to hybrid inviability or behavioral sterility (i.e. hybrids suffer neurological or physiological defects impairing proper courtship behavior) in *Drosophila*^36^. Analyzing epistatic interaction in a haploid system like *Nasonia* males has many advantages. For example, two-loci interactions have significantly fewer genotypes to compare (4 possible genotypes) and there are no interactions (recessive/dominance) between alleles at any particular gene.

## Materials and Methods

### Nasonia stocks

The *Wolbachia*-cured, inbred lines IV7R2 (*N. longicornis* – paternal line) and AsymCHS (*N. vitripennis* – maternal line) were used to produce F_1_ hybrid females. These virgin F_1_ females were isolated as pupae and produced only males due to the haplo-diploid sex determination mechanism in Hymenoptera. F_1_ hybrid females were provided individually with host fly pupae to produce all-male F_2_ offspring (nomenclature used throughout the text = LV[V] with L referring to the *N. longicornis* grandpaternal nuclear genome; V to the *N. vitripennis* maternal nuclear genome; and [V] to the *N. vitripennis* cytoplasm that is maternally derived). Two independent sets of hybrid males were investigated (1^st^ population = 320 and 2^nd^ population = 112 individuals). The observations and genetic analyses of these two populations were performed approximately 5 years apart, by different investigators and in different laboratories (1^st^ population was studied at the University of Würzburg as part of the Dissertation of Christof Pietsch and the 2^nd^ population was investigated at the University of Groningen as part of the Master’s thesis of Sylvia Gerritsma). Behavioral observations on males of the two parental species (140 *N. vitripennis* and 140 *N. longicornis* males) were used for comparison.

### Observations and Test Procedures

Wasps were bred in incubators at 20-25°C under constant light. Test animals (i.e. F_2_ LV[V] males and AsymCHS females) were of controlled age (2-3 days post-emergence). To obtain standardized material males were isolated in glass tubes (75 mm x 12 mm) at least 12 hours prior to testing^37^. All F_2_ hybrid Males were inexperienced whereas females were already mated to prevent premature termination of courtship due to copulation (mated females will not remate, allowing courtship to continue uninterrupted). All males were tested against *N. vitripennis* females to prevent possible effects of female species, although most recorded components of male courtship are not affected by partner species. Neither are they affected by the mating status of the female, since *Nasonia* males seem to be incapable of distinguishing between mated and virgin females^21^. The exception is “total number of cycles” because unmated females allow mating usually after the first 1-4 cycles and non-hybrid males usually give up courtship after about 5-8 cycles.

*Nasonia* male courtship is characterized by a periodically repeated series of motor patterns^11,21,38^. Males approach females with their antennae and if they deem females a suitable partner they climb onto the females back with their legs on the head and thorax of the female. Once the male is in position and the female stops moving, the male starts the courtship behavior with a series of headnods. Successive headnod series are separated by pauses. The interval between the first headnods of two consecutive series is termed a cycle. We recorded in chronological order (1) the time of the males’ rapprochement toward the female called “latency” (interval between introduction of the female and moment of mounting), (2) a character called “fix-nod”, which is the time after the female is immobilized by the male (after arrival at the frontal courtship position) and the onset of the courtship display, (3) the “number of head nods in each cycle” (headnod 1, headnod 2, headnod 3, headnod 4), (4) the “duration of cycle time” (cycle 1, cycle 2, cycle 3, cycle 4) for four consecutive cycles, and (5) the “total number of cycles” until dismounting. *N. longicornis* males show two additional species specific qualitative traits: alternated rubbing with the fore tarsi over the females’ eyes (= “feet rubbing”) and irregularly performed nods without mouthpart extrusions in between the head nod series (=“minus nods”). *N. vitripennis* is characterized by a decline in the number of head nods in the second series, termed “h2-h1”^11,12^. Note that not all behaviors were scored for each individual in both mapping populations.

Observations were made under a dissection microscope at 12 x magnification. Polystyrol tubes (75 mm x 12 mm ∅) served as observation chambers. Observations started with the introduction of a female into the tube and ended with either the male dismounting after a bout of courtship, or after 10 minutes if a male did not mount the female. Males that did not show courtship within 10 minutes were discarded. The software package THE OBSERVER 2.0 (Noldus Information Technology, Wageningen, Netherlands) was used to record the timing of courtship components in the display by tapping specifically coded keys for each component. The second mapping population was observed under the same conditions but instead of using THE OBSERVER the behavior was filmed (Microscope: Leica MZ 12_5_, camera: Diagonostic Instruments Inc. 11.2 Color Mosaic) and scored directly and a second time using the video reording. These video protocols generated an exact time course for every courtship behavior and allowed us to assign time and frequencies to each subcomponent of a male’s courtship behavior.

### Marker amplification

After observation, wasps were stored in 70% ethanol at −20°C. DNA was extracted from whole wasps with a standard phenol-chloroform procedure^39^. Dried DNA was dissolved in 60 μl 0.1 X TE and for PCR the stock DNA was diluted to 5-10 ng/μl. PCR amplification for microsatellite markers was carried out according to Pietsch *et al*.^40^ and Beukeboom *et al.*^41^. PCR products were visualized either with a DNA Analyzer 4300 from LI-COR (1^st^ population in Würzburg) or with an ABI 3730 automated DNA sequencer from Applied Biosystems (2^nd^ population in Groningen). Fragment sizes were determined using GeneScan computer software provided by the manufacturer on the ABI 3730 (Applied Biosystems) or were scored visually when using the LI-COR DNA Analyzer 4300.

### Linkage map construction

A linkage map was built from the genotypes of 320 hybrid males of the first mapping population with a set of 61 microsatellite markers^41^. The second mapping population used 112 males and a subset of 29 evenly spaced microsatellite markers shared with the first mapping population. The software MultiPoint 1.2 (http://www.mulitqtl.com) was used for linkage map construction. A recombination fraction (rf) of 0.3 returned five linkage groups corresponding to the five chromosomes of *Nasonia*. The multilocus ordering was performed with the standard settings in Multipoint (10 iterations, 90% of the population is included in the jackknife analysis). Furthermore the locus ordering was controlled for monotony in order to obtain a reliable framework map for QTL analysis. Linkage groups were assigned to specific chromosomes using chromosome specific markers and the published genome sequence^42,43^. Both linkage maps did not significantly differ in recombination frequency or marker order and cM were calculated using the Kosambi mapping function. Previous comparisons between intra- and interspecific maps found no significant difference in recombination and ordering of markers^41^.

### QTL analysis

MapQTL was used for all interval QTL analyses^44^. QTL were only reported if they passed the 1% genome wide significance threshold determined by running 1000 permutations^45^ the range of the genome wide significance thresholds at 1% for all traits was LOD 2.5-2.8. The R/qtl add-on package for the freely available statistical software R (http://www.r-project.org/) was employed for all interaction analyses^46^. For this analysis, we first conducted standard interval mapping using the scanone-function with the normal model for most courtship components except for minus nods and forefeet rubbing where the binary-model was applied that uses a logistic-regression approach (http://www.rqtl.org/manual/qtl-manual.pdf). 5% genome-wide significance thresholds were then obtained by a permutation test running 1000 permutations^44^. In a second step we ran a two-dimensional QTL analysis using the scantwo-function allowing the estimation of additive and epistatic effects as well as the joint effect of two interacting loci by evaluating four types of models, i.e (1) the full model of additive and epistatic effects, (2) an additive model only, (3) an epistatic model only, and (4) both single QTL models. The two-dimensional scan was performed with a marker-regression approach, after missing genotypes have been filled in by a single imputation. Empirically derived genome-wide thresholds were likewise calculated for a joint QTL model comprising additive and epistatic effects. Significant QTL derived from the two-dimensional genome scan were further analyzed using mean and standard error (SE) of all four haplotypes (VV, VL, LV, LL; V= *N. vitripennis* allele, L= *N. longicornis* allele). In a third step we set up a multiple QTL model according to the following rules: (1) QTL that coincide in a 15 cM interval are treated as a single locus, (2) QTL showing non-significant effects in the single QTL model are still incorporated in the multiple QTL model, if they appear in at least two-thirds of the two-way QTL per trait. The multiple QTL model was likewise calculated with marker-regression. Finally, the model was evaluated by comparing the full model with a reduced model that had a single QTL or epistatic interaction removed at a time. Only QTL that showed a significant reduction (p ≤ 0.05) in the model fit after removal were incorporated in the final model.

For phenotypes for which we found a number of overlapping two-way QTL we performed a multiple QTL scan in a range of ± 15 cM around the QTL positions using the scanqtl-function. Afterward we fitted the QTL-model (with the fitqtl-function) with QTL that decreased the LOD of the entire model below 3 when dropped from the model formula. We only allowed for two-way interactions in our model.

### Testing the effect of sample size on estimated QTL effect

In order to estimate the effect of sample size on the detection and estimation of QTL effects, we performed repeated two-dimensional QTL analysis on different sample sizes for primary focus QTL (see table 2). For that purpose we randomly sampled 100 times (with replacement) sample sizes of 100, 150, 200, 250, 300 and 350 individuals, respectively from the data matrix of our first mapping population. Simultaneously, we determined the 5% genome-wide significance threshold by 1000 permutations for each bootstrap run separately.

### Candidate gene selection

A list of 41 genes involved in courtship behavior (14 genes) and circadian rhythm (27 genes), constructed from *D. melanogaster*^47^, was used as a basis for selecting candidate genes for *Nasonia vitripennis*. NasoniaBase^48^ was used to find the *Nasonia* orthologs of these selected genes and their position in the *Nasonia* genome. Candidate genes were plotted on the linkage map along with the QTL and marker clusters. Confidence intervals for QTL were determined using the drop ≥ 1.5 LOD method^49^ (Table 2). We assigned candidate genes for each QTL if one of the courtship behavior or circadian rhythm genes fell within these confidence intervals. All candidate genes reported were less than 10 cM apart from a significant QTL.

## Results and Discussion

### Phenotypic variations in the F_2_ hybrid male population, transgressive phenotypes and the grandfather effect

Under our experimental paradigm, a large proportion of hybrid males did not show courtship behavior corroborating previous results^12,13^ and reflecting the strong postzygotic isolation between these two species. For that reason about 70% of the tested males had to be excluded from our analyses. This approach made sure that we only scored males that showed a complete courtship behavior and that our results were not biased by males that for example terminated courtship early which would bias the quantification of traits like headnods or cycle time towards smaller values. These unsuccessful males were either not able to get into a proper courtship position on top of the female or did not approach the female in the right way within the 10 minute period they had access to a female. This corroborates previous results using hybrid F_2_ males between *N. vitripennis* x *N. longicornis* or *N. vitripennis* x *N. giraulti*^12,50^. Complete phenotypic data for the QTL analyses were obtained for 320 and 112 individuals of the first and second mapping populations, respectively. Two types of courtship components were recorded, time elements and frequency elements (Figure 1 A, B and C, respectively). Note, no latency times were recorded for the pure species because latency in non-hybrids is typically extremely short (i.e. in the range of a few seconds) whereas latency in hybrids is usually 10-100 times longer (mean = 100 seconds, Figure 1a). Additionally, the variance of every trait in the hybrid mapping population are much higher than in the parental species, which further supports the observation that the hybrid males are phenotypically much more variable than males of either parental species (Table 3).

**Figure 1.**
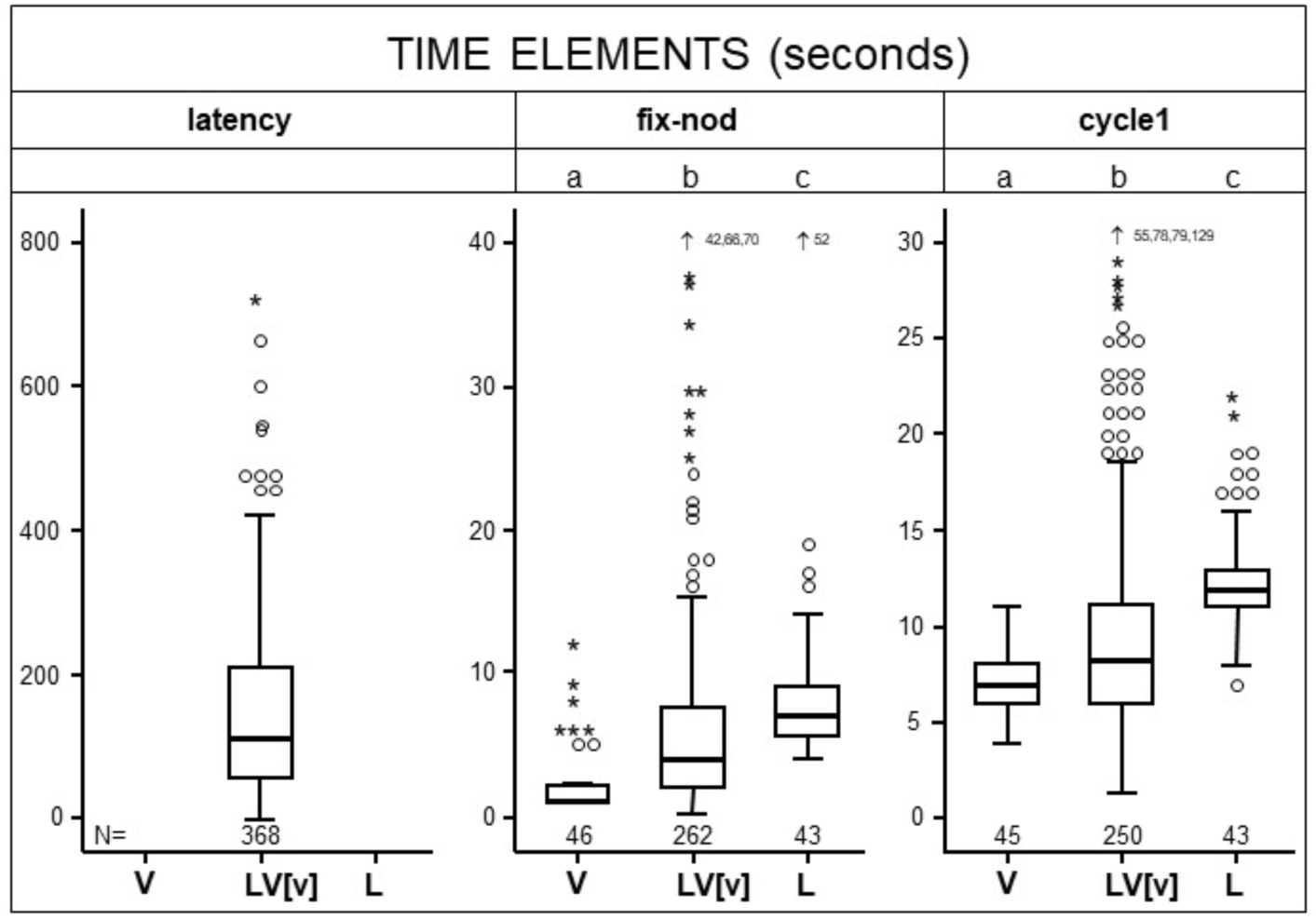
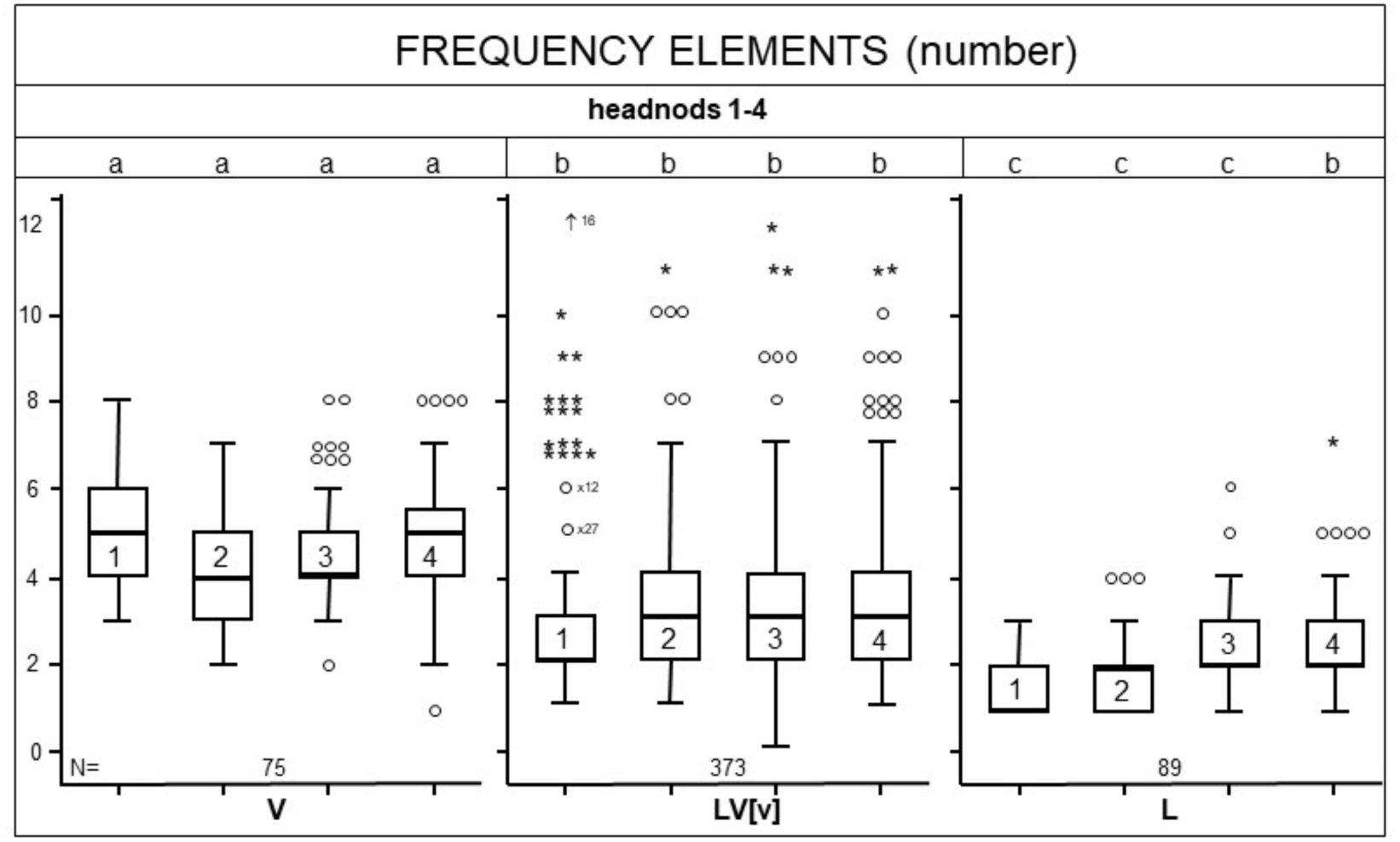
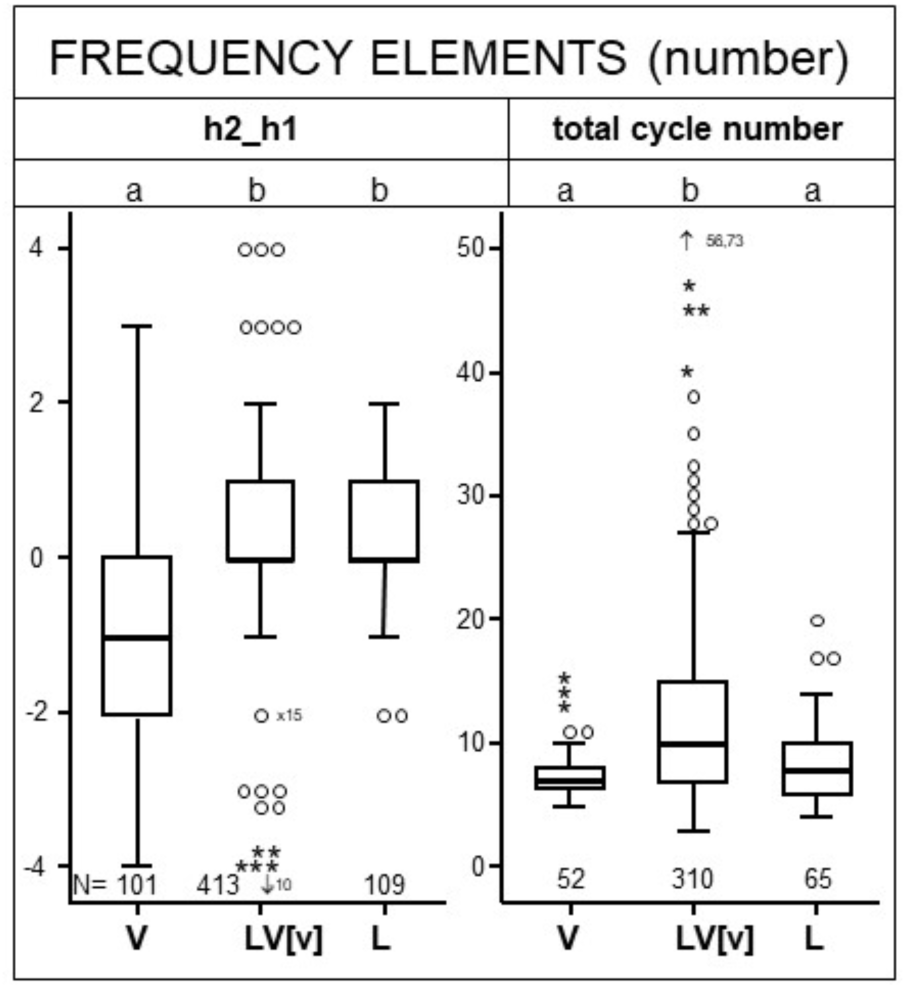
Phenotypic analysis of male courtship behavior of *N. vitripennis* – *N. longicornis* hybrids. (A) Time elements and (B, C) frequency elements. Time elements include duration of latency, fix-nod and cycle 1. Frequency elements include number of headnods in cycles 1-4, h2-h1 and total cycles. For each behavioral component indicated above the graph data are given for both parental species (V = *N. vitripennis*, L = *N. longicornis*) and their hybrid (LV = *N. longicornis* male x *N. vitripennis* female). Data for the hybrids are pooled for the two mapping populations. Box plots show the median (thick horizontal line within the box), the 25^th^ and 75^th^ percentiles (box) and 1.5 times the interquartile range (thin horizontal lines). Outliers are indicated by an open circle and extreme cases are marked by an asterisk. Grouped symbols indicate similar values and arrows with numbers indicate additional extremes that fall beyond the vertical axis ranges. Letters above the graphs indicate significant differences between groups for each behavioral component.

Many of the courtship components are significantly correlated (SOM Table 1). This is not surprising for traits that occur in series (e.g. cycle times 1-4 or headnod series 1-4). In general, all frequency elements and all time elements are strongly correlated among themselves (SOM table 1). Only four traits, cycle time and headnod series as well as cycle times and total number of headnod series (= number of cycles) showed a significant correlation between the two element types. These phenotypic correlations most likely have a genetic basis because some of the QTL for the correlated traits are closely linked (e.g. QTL for fix-nod, cycle1-4 and latency on chromosome 1 (Figure 2)) and most serial traits (e.g. cycle 1-4) share the same QTL (Figure 2).

**Figure 2.**
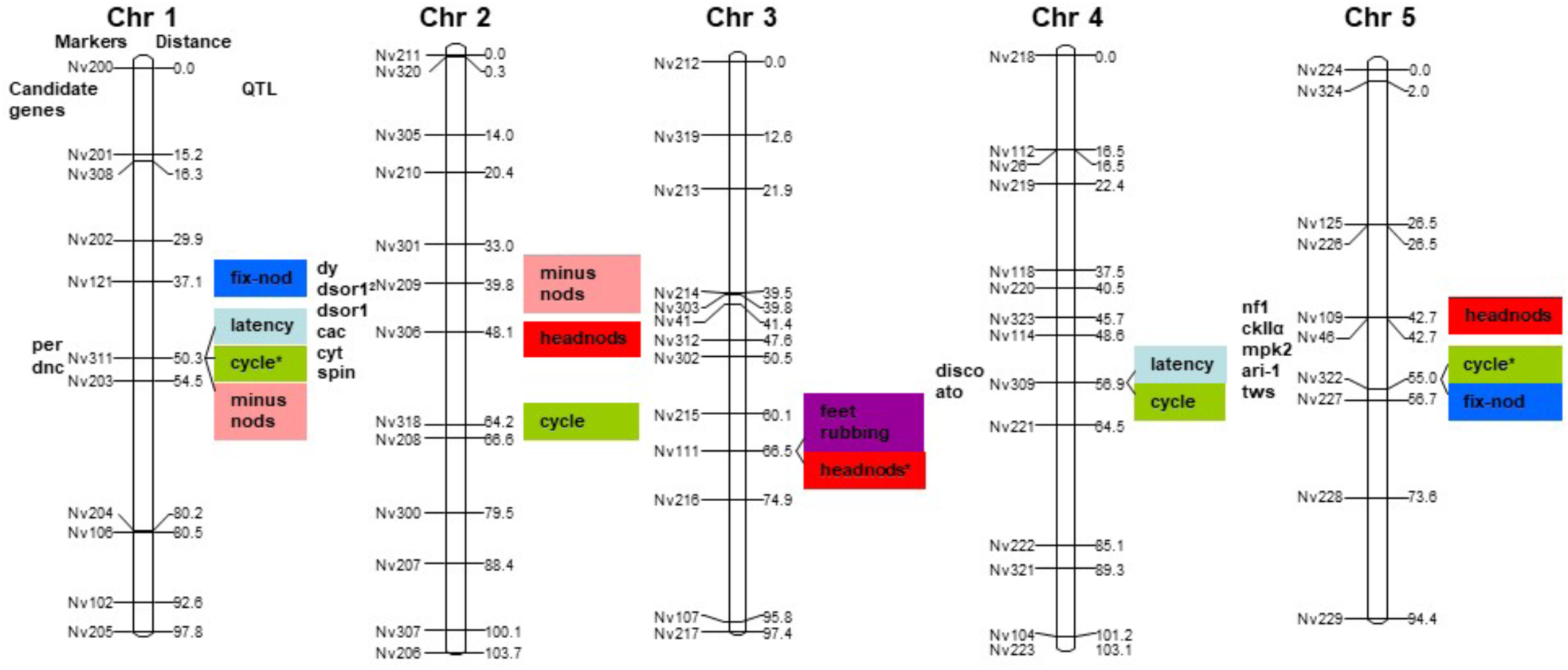
QTL for male courtship behavior of *N. vitripennis* – *N. longicornis* hybrids The linkage map consists of 5 chromosomes. Markers are shown on the left and recombination distance in centiMorgans (Kosambi) on the right of each chromosome. QTL for each trait are color coded and shown on the right side of the closest linked marker. Candidate genes within the confidence interval (1.5≥ LOD drop) of each QTL are shown on left. For their exact location see Table 4. QTL labeled with an asterisk were found in both mapping population. The two QTL headnods and cycle that are marked with an asterisk show significant QTL for headnod 1-4 or cycle 1-4, respectively (see Table 2).

**Table 1.** Association in hybrids between feet rubbing and minus nods. These are unique behaviors that are only performed by *N. longicornis*. The association is significant (χ^2^=7.63, p<0.01).

|  | Feet rubbing | present | absent |
| --- | --- | --- | --- |
| Minus nods |  |  |  |
| present |  | 37 | 50 |
| absent |  | 41 | 120 |

**Table 2.** Significant QTL (p<0.01) for courtship behavior in *N. vitripennis – N. longicornis* F_2_ hybrid males. Data are for two mapping populations; a 1^st^ population of 320 individuals and a 2^nd^ population of 112 individuals. QTL labeled with an asterisk were confirmed in the second mapping population at a minimum genome wide significance thresholds of 1%. LOD scores for the QTL in the second mapping population are given in italics in the fourth column (LOD). The explained phenotypic variance for the QTL in the second population is shown in italics after the explained phenotypic variance for the same trait and locus in the larger population. Gray backgrounds mean loci (column chr.=chromosome) with bootstrapping estimates for phenotypic effect of both *LOD* scores and *explained variance* (see figure 1, Supplementary Online Material) and loci (column mean long allele and mean vit allele) with an inverse effect based on the parental phenotype, i.e. *N. vitripennis* shows more headnods than *N. longicornis* but the substitution of a *N. vitripennis* allele at chromosome III reduces the average number of headnods in the hybrid male mapping population. The columns mean long allele and mean vit allele show the mean phenotypic value for F_2_ hybrid males that inherited a *N. vitripennis* or *N. longicornis* allele at the marker most closely associated with the QTL.

| trait | chr. | position [cM] | LOD | Drop $\geq 1.5$ LOD interval (LOD score at flanking marker) | explained variance [%] | mean long allele | mean vit allele | marker |
| --- | --- | --- | --- | --- | --- | --- | --- | --- |
| latency | I | 50.3 | 4.3 | 37.1-64.5 (2.5-2.8) | 7.2 | 187.7 | 123.9 | Nv311 |
|  | IV | 56.9 | 3.8 | 40.5-74.5 (1.8-2.1) | 6.2 | 186.6 | 125.7 | Nv309 |
| fix-nod | I | 37.1 | 4.1 | 21.3-59.5 (2.5-2.5) | 6.7 | 9.46 | 4.83 | Nv121 |
|  | V | 56.7 | 4.4 | 42.7-78.6 (2.9-2.7) | 7.1 | 9.63 | 4.83 | Nv227 |
| total series | V | 55 | 29 | 36.5-66.7 (1.2-0.8) | 6.4 | 9.5 | 14.2 | Nv322 |
| cycle1 | I* | 50.3 | 7.1/<br>2.6 | 37.1-59.5 (5.0-5.4) | 10.5/15.7 | 15.5 | 7.95 | Nv311 |
|  | II | 64.2 | 5.1 | 48.1-84.5 (3.2-2.6) | 7.6 | 14.01 | 7.90 | Nv318 |
|  | V* | 55 | 5.9/<br>4.9 | 42.7-71.7 (3.8-3.1) | 9.0/25.9 | 14.76 | 7.96 | Nv322 |
| cycle2 | I* | 50.3 | 9.2/<br>2.5 | 37.1-64.5 (7.1-6.7) | 13.5/14.1 | 14.62 | 8.92 | Nv311 |
|  | II | 64.2 | 7.1 | 48.1-79.5 (3.7-5.2) | 10.5 | 13.71 | 8.89 | Nv318 |
|  | IV | 56.9 | 3.1 | 45.7-79.5 (1.6-1.6) | 4.7 | 12.91 | 9.60 | Nv309 |
|  | V* | 55 | 11.5<br>/5.2 | 42.7-56.7 (6.1-9.5) | 17.1/24.5 | 14.9 | 8.60 | Nv322 |
| cycle3 | I* | 50.3 | 9.1/<br>6.6 | 37.1-64.5 (5.1-6.6) | 18.2/32.9 | 13.79 | 8.74 | Nv311 |
|  | II | 66.6 | 3.3 | 84.5-39.8 (1.8-1.2) | 7.1 | 12.22 | 9.19 | Nv208 |
|  | V | 55 | 9.3 | 36.5-61.7 (7.9-7.7) | 19.5/16.2 | 13.96 | 8.71 | NV322 |
| cycle4 | I* | 50.3 | 12.6<br>/4.6 | 37.1-54.5 (8.9-10.3) | 25.5/19.6 | 13.62 | 8.98 | Nv311 |
|  | IV | 56.9 | 3.7 | 32.4-74.5 (2.2-1.8) | 8.2 | 12.09 | 9.53 | Nv309 |
|  | V* | 55 | 8.1/<br>5.7 | 42.7-66.7 (6.3-6.5) | 17.7/27.5 | 13.12 | 9.24 | Nv322 |
| headnod1 | II | 48.1 | 5.0 | 39.8-63.1 (1.7-3.0) | 7.4 | 3.34 | 2.41 | Nv306 |
|  | III* | 66.5 | 5.0/<br>3.5 | 60.1-94.9 (2.1-3.2) | 7.3/13.3 | 2.18 | 3.11 | Nv111 |
|  | V | 42.7 | 3.8 | 7-56.7 (1.9-2.3) | 5.5 | 3.30 | 2.61 | Nv109 |
| headnod2 | III* | 66.5 | 6.4/<br>4.4 | 60.1-74.9 (4.0-4.9) | 9.2/16.3 | 2.30 | 3.31 | Nv111 |
|  | V | 42.7 | 7.3 | 26.5-52.7 (4.6-5.8) | 10.4 | 3.64 | 2.56 | Nv109 |
| headnod3 | III | 66.5 | 4.3 | 60.1-89.9 (2.5-2.1) | 6.3/6.1 | 2.57 | 3.43 | Nv111 |
|  | V | 42.7 | 11.5 | 31.5-55.0 (9.1-9.5) | 16 | 4.01 | 2.63 | Nv109 |
| headnod4 | III* | 66.5 | 3.34<br>/2.7 | 60.1-94.9 (1.8-1.7) | 5.2/9.3 | 2.71 | 3.49 | Nv111 |

**Table 3:** Explained genetic variance (both additive and epistatic) in male courtship behavior using the full QTL model for each trait. Columns 2-4 show the additive, epistatic and total explained phenotypic variance in our maping population, columns 5-6 show the variance of the same traits in the isogenic non-hybrid males from the parental species (var-vit = *N. vitripennis*; var-lon = *N. longicornis*, See SOM table 4), i.e. this variance reflects mostly environmental variance, and columns 7-8 show the phenotypic variance in the large (1st) and small (2nd) F_2_ hybrid male mapping population.

| trait | expl.<br>var.<br>add | expl.<br>var.<br>epi. | expl.<br>var.<br>total | Var-Vit | Var-Lon | Var-LV<br>1st | Var-LV<br>2nd |
| --- | --- | --- | --- | --- | --- | --- | --- |
| latency | 4.97 | 0 | **4.97** | n.a. | n.a. | **12736.67** | **16565.063** |
| fix1nod | 32.06 | 2.20 | **34.26** | 6.047 | 57.245 | **73.953** | **n.a.** |
| cycle1 | 39.61 | 8.71 | **48.31** | 2.77 | 6.38 | **117.69** | **39.207** |
| cycle2 | 33.86 | 2.65 | **36.51** | 1.32 | 3.51 | **52.07** | **29.614** |
| cycle3 | 27.56 | 1.44 | **29.00** | 1.54 | 1.89 | **29.96** | **37.628** |
| cycle4 | 18.94 | 0.00 | **18.94** | n.a. | n.a. | **17.89** | **37.054** |
| headnod1 | 21.63 | 0.00 | **21.63** | 1.672 | 0.638 | **2.774** | **4.089** |
| headnod2 | 29.43 | 0.00 | **29.43** | 1.303 | 0.489 | **2.546** | **2.859** |
| headnod3 | 29.95 | 0.00 | **29.95** | 1.575 | 0.825 | **2.731** | **3.155** |
| headnod4 | 23.63 | 0.00 | **23.63** | 1.677 | 1.001 | **2.781** | **3.421** |
| h2_h1 | 4.77 | 0.00 | **4.77** | 1.45 | 0.48 | **1.46** | **1.761** |
| minusnods | 12.29 | 3.16 | **15.45** | n.a. | n.a. | **0.231** | **n.a.** |
| Feet<br>rubbing | 6.93 | 1.85 | **8.78** | n.a. | n.a. | **0.22** | **n.a.** |

**Table 4.** Candidate genes for courtship behavior QTL in *Nasonia*. Distance reflects the distance in centiMorgans (Kosambi)of the gene to the marker closest associated with the trait in question.

| Marker | Chromosome. | Trait | Gene | Functional basis | Distance (cM) |
| --- | --- | --- | --- | --- | --- |
| Nv121 | 1 | Fix-nod | *Period* | Circadian rhythm | 4.5 |
| Nv311 | 1 | Latency, Cycle1, Cycle2 Cycle3, Cycle4 | *Period* | Circadian rhythm | 3.6 |
|  |  |  | *Dunce* | Circadian rhythm | 2.7 |
| Nv318<br>Nv208 | 2 | Cycle1, Cycle2<br>Cycle3 | *Spinster* | Circadian rhythm | 2.7 |
|  |  |  | *Synaptotagmin* | Circadian rhythm | 0.0 |
|  |  |  | *Cacophony* | Courtship behavior | 7.4 |
|  |  |  | *Dopa decarboxylase* | Courtship behavior | 7.4 |
|  |  |  | *Downstream of raf1* | Circadian rhythm | 8.3 |
|  |  |  | *Dusky* | Circadian rhythm | 9.2 |
| Nv306 | 2 | Hnd1 | *Synaptotagmin* | Circadian rhythm | 7.4 |
|  |  |  | *Cacophony* | Courtship behavior | 0.0 |
|  |  |  | *Dopa decarboxylase* | Courtship behavior | 0.0 |
|  |  |  | *Downstream of raf1* | Circadian rhythm | 0.9 |
|  |  |  | *Dusky* | Circadian rhythm | 1.8 |
| Nv111 | 3 | Hnd1, Hnd2, Hnd3, Hnd4 | - | - | - |
| Nv309 | 4 | Latency, Cycle2, Cycle4 | *Disconnected* | Circadian rhythm | 1.0 |
|  |  |  | *Atonal* | Courtship behavior | 2.8 |
| Nv109 | 5 | Hnd1, Hnd2, Hnd3, Hnd4 | *Neurofibromin 1* | Circadian rhythm | 0.9 |
|  |  |  | *Casein kinase II alpha subunit* | Circadian rhythm | 7.2 |
|  |  |  | *Mpk2* | Circadian rhythm | 7.2 |
|  |  |  | *Ariadne* | Courtship behavior | 9.0 |
| Nv322 | 5 | Cycle1, Cycle2<br>Cycle3, Cycle4 | *Ariadne* | Courtship behavior | 0.0 |

Mean hybrid courtship components are typically intermediate between both parental species with the exception of total number of cycles, which was considerably higher for the hybrids^12,13^(see Table 4 for all parental phenotypes). This is in accordance with previous findings and confirms that most F_2_ hybrid males that initiate courtship are missing the mechanism that non-hybrid males use to stop courtship behavior if the female is not becoming receptive within an appropriate time frame. The females that were used in the courtship experiments were all mated and hence would never become receptive even when courted by non-hybrid males under the experimental paradigm. This courtship-stopping mechanism is probably an adaptive behavior because males under natural conditions will encounter a mixture of mated and unmated females and they will lose mating opportunities if they court an unreceptive female for too long.

The previously reported grandfather effect is also evident in our mapping populations for headnod series 1-4 and the structure of the courtship bout, exemplified by the number of headnods in cycle 2 minus cycle 1 (h2-h1)^12,13^. Hybrids resemble *N. longicornis* more for these traits even though headnod series 1-3 differ significantly in hybrids from both pure species (Figure 1). Our results did not provide any detailed insight into the genetic basis of the grandfather effect but we can exclude epistatic interactions between the identified QTL as an explanation because there was no epistatic interactions between any of our headnod QTL (Table 3).

Some of the F_2_-hybrid males show transgressive phenotypes, i.e. phenotypes that fall significantly outside the range of the parental phenotypes (Figure 1). For example seven out of 320 males show more than 8 head nods in headnod 1, which was never observed in any of the parental species (Figure 1b, SOM table 3). These extreme phenotypes are common in animal and plant hybrids and can be important in the formation of hybrid species because they might allow hybrids to successfully exploit niches significantly different from their parental species^34^. Alternatively, transgressive phenotypes in the case of male courtship behavior can also act as postzygotic isolation mechanisms reducing hybrid fitness. Since our individuals are haploid there are no intra-locus interaction effects (e.g. dominance of one allele), i.e. these observed phenotypes reflect the true phenotypic effect of an allele in any given genetic background (for interactions between two or multiple loci see below). Table 2 (columns 6+7) lists the mean effect of the *N. vitripennis* and *N. longicornis* alleles on the trait under consideration. This allows us to determine whether the phenotypic effect is in the direction of the parental line it originated from, e.g. since the mean cycle time of *N. longicornis* is longer (±12 s, Figure 1) than that of *N. vitripennis* (±8 s,) we would expect that individuals in the F_2_ population that inherited a *N. longicornis* allele at any QTL locus for cycle time should show a higher mean than hybrids inheriting a *N. vitripennis* allele at this locus. This is what we observe for all cycle time QTL but not for headnod QTL (Table 2). In two of the headnod QTL (II-48.1 and V-42.7, table 2) the allelic effect is reversed, i.e. the *N. longicornis* allele adds on average one additional headnod if it is substituted for a *N. vitripennis* allele at this QTL locus (Table 2). Since some of the individuals in the larger population show transgressive phenotypes for headnods (Fig. 1b) it was expected that those individuals should have both *N. longicornis* alleles at QTL II-48.1 and V-42.7 but a *N. vitripennis* allele at QTL III-66.5. This was indeed the case for most individuals that showed transgressive, phenotypes (n = 7). However, other unknown genetic or environmental factors seem to be important because not all individuals with these genotypes showed transgressive phenotypes.

### Genetic architecture of species differences in male courtship behavior

#### Interval Mapping

We first performed standard interval mapping in mapping population 1 (n = 320 F2 hybrid males), revealing 26 significant QTL at 14 independent loci (Table 2, Figure 2). Two to four QTL were found per trait for 10 out of 11 quantitative traits analyzed. Surprisingly, no significant QTL were detected for h2-h1 despite the clear species specific pattern. QTL for headnods 1-4 and cycle 1-4 were mostly co-localized (pleiotropy), i.e. the major QTL effect of the different cycles 1-4 was always associated with the same marker (Figure 2). Ten QTL of the 26 previously identified QTL could be confirmed in a second independent mapping population conducted five years later (QTL marked with an asterisk in Table 2). No additional QTL were found. The reason for not recovering the remaining QTL is most likely the smaller population size that makes it more likely to miss QTL with a smaller phenotypic effect. This explanation is supported by the significantly lower explained variance of missed QTL versus confirmed QTL in the second analysis (Table 2, Mann Whitney-U p=0.028). An alternative explanation would be that the two strains used in the second QTL analysis changed their genetic composition significantly, but we have no evidence for this from other studies using the same strains.

Both the individual QTL effects (explained variance per locus) and the percentage of total variance explained (sum over all loci) varied considerably between traits and loci (Table 2 and 4). For the three traits (total number of cycles, latency and fix-nod) we found only one or two loci each that explained between 6.2 and 7.4 % of the observed phenotypic variance in the first mapping population and neither of those QTL could be confirmed in the second, smaller mapping population. For both serial traits (headnods 1-4 and cycles 1-4) we found multiple independent QTL. The major loci for headnods and cyles on chromosomes I, III and V respectively, were confirmed in the second smaller mapping population. QTL for headnods 1-4 or cycles 1-4 explained between 5.2 and 25.5 %, and 6.1 and 32.9% of the observed variance in the first and second mapping population, respectively. These differences in explained effect size can be attributed to the so-called Beavis effect and we deem the values of the larger mapping population more reliable as they also come closer to the means in the permutation tests (Figure 4, see also discussion on the Beavis effect below).

It is interesting that the percentage of explained variance increases for the two confirmed QTL loci within the cycle series (cycle 1-4 QTL on chromosome I increases from 10.5% to 25.5%; cycle 1-4 on chromosome V from 9.0% to 17.7%, Table 2), but not in the headnod series. Whether this reflects a difference in the underlying genetic architecture or a statistical artifact is unclear. Interestingly, the number of two-way interactions increase from cycle 1 (n=10) to cycle 4 (n=14) whereas they decrease from headnod 1 (n=12) to headnod 4 (n=7) (Figure 3b and 3c). This and the fact that no non-additive epistatic interactions were found (SOM Table 2) for headnods might explain the differences in explained variance over consecutive series of cycle time and headnods. Our study finds both QTL with large and small effects and therefore does not support the notion that the genetic architecture underlying species differences in prezygotic isolation mechanisms generally involves few loci with major effect^2^. Rather our results support the view that limited/small sample sizes lead to a bias towards QTL with large effect due to the reduced power to detect QTL with small effect (small mapping population = 10 QTL *versus* large population = 26 QTL, see also figure 4 – distribution of QTL effects with smaller sample sizes range below significance threshold). Our results are support a mixed model where both QTL with small and large effect contribute to the observed species differences.

**Figure 3.**
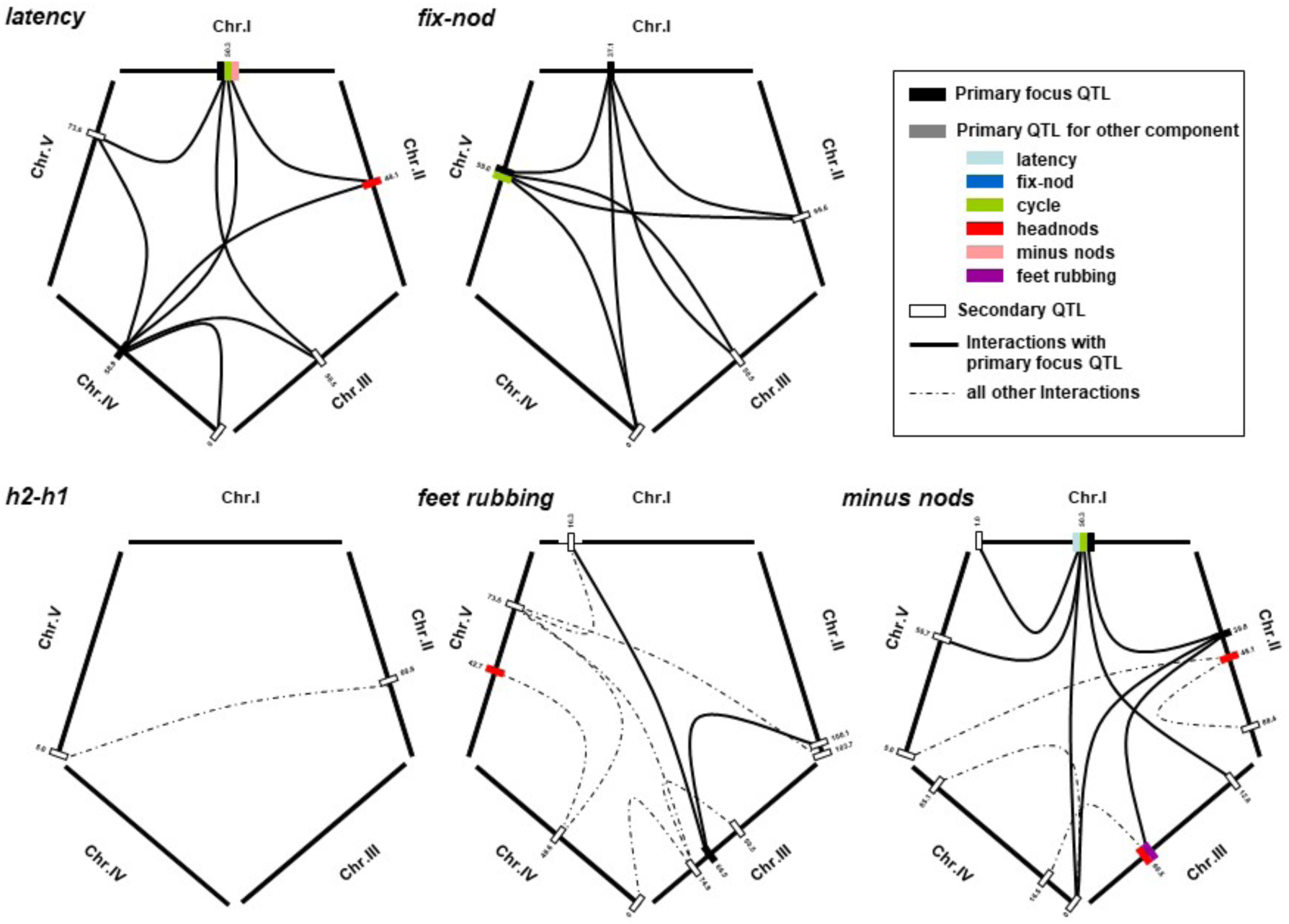
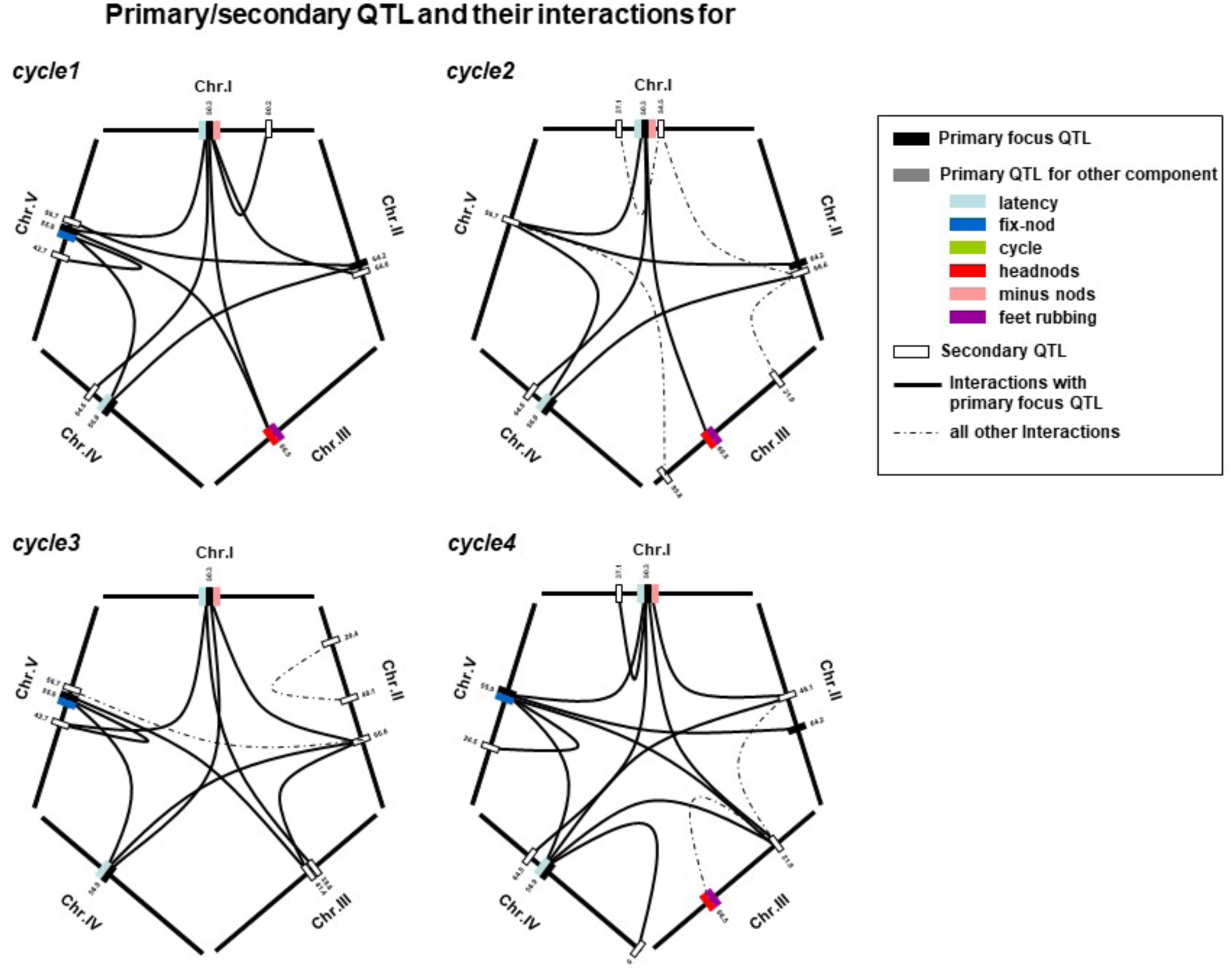
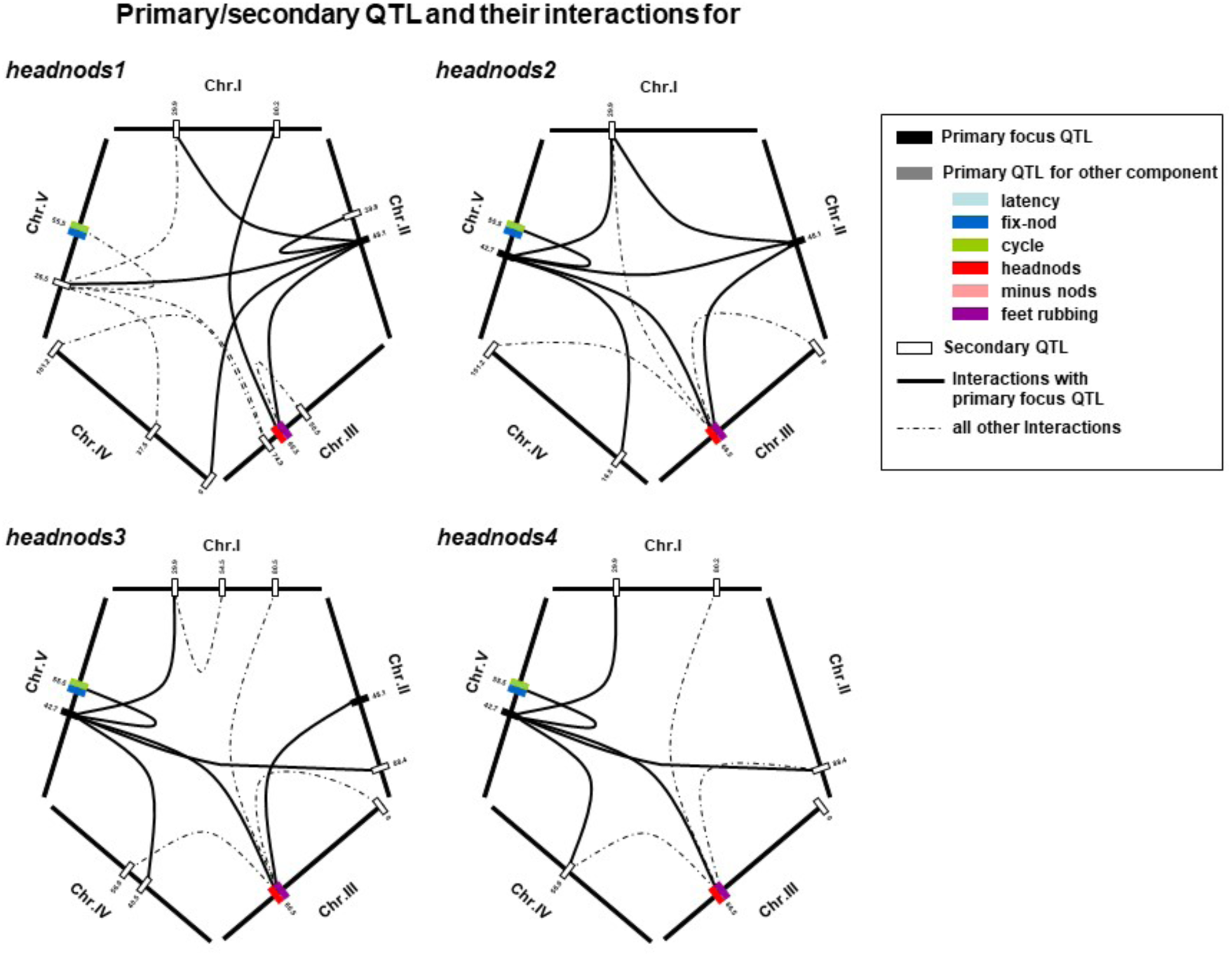
Significant two-way interactions between male courtship QTL. Significant epistatic interactions are shown for each courtship component. The courtship component under focus is indicated at the upper left corner of a pentagon. Each side of the pentagon represents one of the five chromosomes of *Nasonia*. Thick lines indicate epistasis between a focus trait QTL and a secondary QTL, thin lines connect secondary QTL with either QTL for other traits or other secondary QTL. Colored rectangles on the chromosomes correspond to the QTL in figure 2, the focus QTL are shown by black rectangles and the secondary QTL with white rectangles. Note that repeated components (e.g. cycles 1-3, head nods 1-4) are strongly correlated and show similar but not identical epistasis patterns.

#### Two Way interactions and complete Models

The basic assumption of quantitative genetics is that the genetic architecture underlying variation in quantitative traits is polygenic and all/most loci have small and additive phenotypic effects^50^. Most studies on the genetic basis of species differences or differences in insect courtship behavior found, in contrast, few genes and a majority of those had major phenotypic effects (table 6.2 in ^1,2,51^). The results of our study do not corroborate these results (see above). Additionally, since we worked with haploid individuals we are in a much stronger position to investigate the contribution of interacting nuclear loci (additive or epistatic) on the observed phenotypic variance. Hence, our results provide information on the validity of the assumption that the genetic basis of quantitative traits interact additively. For that purpose, we conducted a whole genome two-way interaction analysis in which we searched for loci that had no significant effect on the phenotypes on their own (Fig. 3 secondary loci) but had a significant phenotypic effect in connection with another locus. The result of this analysis is summarized in the SOM (Table 2) and the individual interactions are graphically depicted in Figure 3. These interacting loci were either modifiers of large effect QTL (e.g. Fig. 3, interaction between primary focus QTL and secondary loci in cycle 1-4) or loci whose effect is conditional on the genetic background at another locus, that also has no effect on the phenotypic variance on its own (Fig. 3, headnods 1 interaction between two secondary loci). Most of the significant interactions were between focal QTL and secondary loci, i.e. one or multiple loci modified the effect of a major QTL (Figure 3). This is in accordance with ideas about the evolutionary dynamics of novel mutations. Large effect mutations rarely have positive fitness effects but even if they do they will often have negative pleiotropic side effects (e.g. ^52^). If this new mutation/allele increases in frequency selection will favor the evolution/increase in frequency of modifier alleles at other loci that ameliorate the negative side effects of this new allele. If we apply this concept to our results this would suggest that the major phenotypic differences in the male courtship behavior between *N. vitripennis* and *N. longicornis* is based on few genes with large effects (QTL in Table 2, primary focus QTL in Figure 3) whose initially negative pleiotropic side effects have later been modified by many unlinked nuclear loci that have no phenotypic effect of their own (secondary QTL in Figure 3). Stern and Orgogozo^42^ and others have argued that epistasis reduces the rate of evolution because selection can only act on the effect of this allele in certain genetic backgrounds. Hence long term selection should favor nonepistatic alleles. From our results we cannot determine whether the large number of interacting loci is due to hybridization or reflects the true genetic architecture of courtship behavior within species. However, it indicates rapid evolution of modifying loci after the first QTL with large effect originally arose in the ancestral population of the two species.

The co-localization of interacting loci for one trait with primary QTL for other traits might indicate a more complex form of pleiotropy (Figure 3), namely that QTL that have a significant direct effect on one trait also have an indirect effect as secondary QTL for another trait (= indirect pleiotropy). We also tested whether those two way interactions were epistatic or additive^33^. Previous studies on morphological traits (wing size and head size) in hybrid males of *Nasonia* revealed many epistatic and non-additive interactions^28^. However, to our surprise most of the interaction for the male courtship behavior traits were additive (>90%, Table 1 in Appendix (two-way interactions)) and non-additive epistatic interaction explained only a small proportion of the observed variance (0 - 8.71%, Table 4). Hence, the observed transgressive phenotypes cannot be attributed to epistatic interactions or more precisely epistatic two way interactions of the QTL we identified.

#### Genetic basis of qualitative species differences in male courtship behavior

Feet rubbing and minus nods, behaviors that are unique to *N. longicornis*, showed a significant association within the F_2_ males population (χ^2^=7.63, p<0.01, Table 1) which indicates either close linkage between the responsible genes or pleiotropic effects. To find the pleiotropic locus or loci that explain some of the variance of these two binomial traits, we conducted chi-square tests at each of the 61 markers of the 320 individuals of the first mapping population. After Bonferroni correction, one locus showed a significant association with feet rubbing (chromosome III *χ^2^* = 16.93, n=320) and two loci a significant association with minus nods (chromosome I (*χ^2^* =23.35) and II (*χ^2^* = 20.92). Note that these traits were not scored in the second population. However, none of the three QTL for those two traits are on the same chromosome (Fig. 2). Hence, we can dismiss pleiotropy for the observed phenotypic correlation between feet rubbing and minus nods in our mapping population unless we missed a major pleiotropic QTL. Another explanation would be two-way interactions between unlinked loci and indeed we found a strong two-way interaction between one of the two QTL for minus nods (Chr.2-39.8) and the only primary QTL for feet rubbing (Chr. 3-66.5) (Fig. 3a).

### Effect of sample size on detection and estimation of explained variance (Beavis effect)

QTL effects are often overestimated owing to the inflation of the effect size by sampling errors^7–9^. Sampling error in this context means that it is more likely to detect a QTL that by chance seems to explain a higher variance than it really does because the power to detect QTL is linked with the size of its effect, i.e. if the explained phenotypic effect is too small the QTL is not significant and can not be reported. The power to detect QTL with small effects is also increasing with sample size^8^. This can be seen in Figure 4 for cycle 1 where at a sample size of 100 males roughly half of the time (50 out of 100 resampled populations) the QTL for cycle 1 at position I-50.3 does not pass the LOD significance threshold (LOD 2.7 = 1 % genome wide confidence interval) and the ones that do pass overestimate on average the explained variance in the full mapping population of 320 males. Note that explained variance and LOD score are highly correlated (in our simulations correlation coefficients are always higher than 0.99). The fraction of significant QTL tends therefore to be biased for QTL with large effects especially if mapping populations are small, which is known as the Beavis effect^8^. In order to assess the effect of sample size on the overestimation of QTL effects in our own data and to determine whether we are overestimating our QTL effects, we performed a bootstrap analysis for the largest QTL for continuous and discrete courtship components. The distribution of estimated QTL effects over sample size showed a similar progression irrespective of the discrete or continuous nature of the courtship components and the differences in the LOD scores ranging from 4,83 for the courtship component total number of cycles to 27,39 for cycle 2 in the initial two-dimensional QTL analysis (Figure 3).

**Figure 4.**
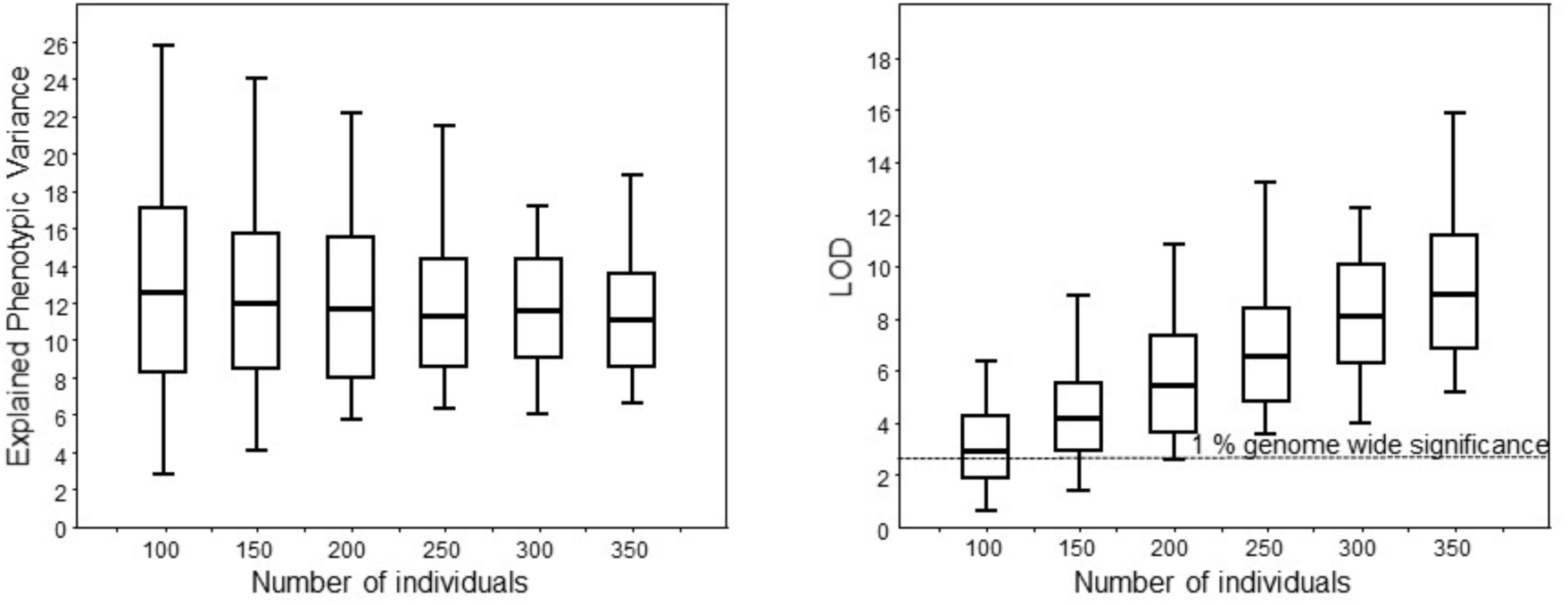
Effect of sample size on estimation of explained phenotypic variance (Beavis effect) and LOD score for the cycle 1. Explained phenotypic variance for cycle 1 is shown as a function of sample size based on a bootstrap analysis (1000 bootstraps/sample size). Boxplots show means/SD/min-max. In a significant number of permutations, QTL would go undetected because they fall below the genome wide detection limit of 1% if the number of individuals in a mapping population is lower than 200 (for most traits the 0.01 significance threshold is around LOD 2.7, dashed line in the right panel). As predicted, the variance in the estimation of a QTL effect becomes smaller as sample size goes up and significant QTL in our studies should always be detected if mapping populations are larger than 250 individuals (all LOD scores are above the detection threshold, left panel). Only results for one QTL (Cycle 1) are shown because the effect is similar for all other courtship components (see SOM Figure 1 for the results for the other traits).

The largest differences of the estimated QTL effects occurred in our data sets between a sample size of 50 and 100. Once a sample size of 250 was reached, changes in estimated QTL effects became considerably smaller. The overestimation of mean QTL effects with a sample size of 50 compared to the mean effects with n =350 range from a 2.57 fold overestimation for headnod 2 to a 1.46 fold overestimation for cycle 4 (Figure 4). Since the sample sizes on which we based our estimates of the QTL effect for all of our QTL analyses were all larger than 300 we consider the estimated QTL effects of this study as accurate. Our results confirm the importance of using sufficiently large sample sizes in QTL analyses ^52^.

### Candidate genes

We identified orthologous genes in *Nasonia* that are involved in courtship behavior and circadian rhythm in *Drosophila melanogaster*. These potential candidate genes were then mapped and checked whether they fall within the confidence intervals of any of our QTL (drop ≥ LOD 1.5 interval, Table 2). This approach produced 14 candidate genes (Figure 2, Table 3). The candidate courtship behavior genes have functions that are involved in various biological processes like localization, signal transmission, mating, locomotory behavior (*cacophony*), catecholamine neurotransmission (*Dopa decarboxylase*), anatomical structure development (*atonal*), growth control, signal transduction, and viral pathogenicity (*ariadne*). The remaining genes are listed as circadian rhythm genes and may be involved in biological processes such as circadian behavior (*period*), locomotor rhythm (*disconnected*, *Casein kinase II α subunit*), locomotory behavior (*spinster*), neurotransmission (*synaptotagmin*), responses to extracellular signals, such as hormones, light, and neurotransmitters (*dunce*), intracellular signaling pathways (*downstream of raf1*, *dusky*), and response to stress (*Mpk2*, *Neurofibromin*)^53^. This result is encouraging and may hind at evolutionary conservation or evolutionary predictability sensu Stern and Orgogozo^41^ of target loci for modifications of male courtship behavior in insects. In the future we will test whether expression of these genes differ systematically between males of the two species and whether we can correlate differences in their expression to differences in the courtship behavior components of F_2_ hybrid males. Additionally, since dsRNAi is working in *Nasonia* we can also analyze the effect of candidate gene knockdowns within both species.

## Conclusions

Knowledge of the genetic architecture of courtship behavior is still rudimentary. The review by Arbuthnott^2^ came up with very few studies and those few studies indicate that most reported courtship traits are regulated by few loci with large effects. However, these studies are strongly biased towards *Drosophila* and in many cases sample sizes may not have been sufficient to detect QTL with small effect. Our comparison of two independent QTL studies that differ significantly in sample size reaffirms this notion because QTL with smaller effects were only detected in the larger population. But, we were able to confirm 10 major QTL in this independent study with a much smaller mapping population. Hence, there is a tradeoff between the number of QTL detected and sample size and it depends on the expected QTL effect size. QTL with large effect can be assessed with significantly smaller sample sizes (50 instead of 300 males) than QTL with small effect. The ability to identify candidate genes for all of the confirmed QTL loci should make it possible to follow up and identify the underlying gene/s using eQTL studies in hybrid populations or dsRNAi knockdown studies in the parental species^54^.

Another unanswered question is whether the discerned genetic architecture of courtship behavior differs when using interspecific versus intraspecific crosses^55^. To resolve this we would need QTL studies on behavioral variants within species which may be rare in nature.

We found extensive epistatic interactions between different behavioral components. This could reflect the genuine genetic architecture of courtship, but it might also be due to the haplodiploid mode of reproduction of *Nasonia*. At this point we do not know whether haploidy of males facilitates the evolution of gene interactions or merely increases the power of detecting epistasis. Resolving this requires further investigation of the genetic architecture of traits in haplodiploids as well as more detailed studies of courtship behavior in diploids.

This study is a major step forward in identifying genes involved in behavioral differences between species in general and reveals some aspects of the genetic architecture of prezygotic isolation mechanisms in *Nasonia* in particular. Once the loci (genes or regulatory units) that form the basis of species differences in male courtship behavior in *Nasonia* spp. have been identified we will have a better understanding of the genetic changes that have led to the evolution of prezygotic isolation in *Nasonia*. We know that chemical communication/signals play an important role during courtship behavior and have already started work on the genetic basis for those signals^56–58^. Our ultimate goal is to gain a full understanding of all genetic factors involved in prezygotic isolation in the genus *Nasonia*.

## Supporting information

SOM Table1+2+3

SOM Figure 1

## Acknowledgements

We thank two reviewers for helpful comments that improved the manuscript and Joshua Gibson for improving the readability of the manuscript. LWB was supported by Pioneer grant no. 833.02.003 from the Netherlands Organisation for Scientific Research. JG was supported by NIH R24 GM084917 and SFB 554 from the Deutsche Forschungsgemeinschaft. We also acknowledge the contribution of two colleagues J. van den Assem and S. Ferber who unfortunately passed away.

## References

1. Coyne JA, Orr HA (2004) Speciation. 545 pp, Sinauer Ass., Sunderland Mass.

2. Arbuthnott D (2009) The genetic architecture of insect courtship behavior and premating isolation. Heredity 103: 15–22.

3. Coyne JA, Orr HA (1989) Patterns of speciation in Drosophila. Evolution 43: 362–381.

4. Coyne JA, Orr HA (1997) “Patterns of speciation in Drosophila” Revisited. Evolution, 51: 295–303.

5. Jaenike J, Dyer KA, Cornish C, Minhas MS (2006) Asymmetrical Reinforcement and *Wolbachia* Infection in Drosophila. PLoS Biol 4(10): e325. doi:10.1371/journal.pbio.0040325

6. Mallet J (2008) Hybridization, ecological races and the nature of species: empirical evidence for the ease of speciation. Phil Trans R Soc B 363: 2971–2986.

7. Beavis WD (1994) The power and deceit of QTL experiments: lessons from comparitive QTL studies, pp. 250–266 In: Proceedings of the Forty-Ninth corn & Sorghum Industry Research Conference. American Seed Trade Association, Washington, DC.

8. Beavis WD (1998) QTL analyses: power, precision, and accuracy, pp. 145–162 In: A. H. Paterson. Molecular Dissection of Complex Traits. CRC Press, New York

9. Xu S (2003) Theoretical Basis of the Beavis Effect Genetics. 165:2259–2268

10. Rockman MV (2012) The QTN program and the alleles that matter for evolution: all that’s gold does not glitter. Evolution 66: 1–17.

11. van den Assem J, Werren JH (1994) A comparison of the courtship and mating behaviour of three species of *Nasonia* (Hymenoptera: Pteromalidae). J Insect Behav 7: 53–66.

12. Beukeboom LW, van den Assem J (2001) Courtship and mating behaviour of interspecific *Nasonia* hybrids (Hymenoptera, Pteromalidae): a grandfather effect. Behav Genet 31: 167–177.

13. Beukeboom LW, van den Assem J (2002) Courtship displays of introgressed interspecific hybrid *Nasonia* males: further investigations into the ‘grandfather effect’. Behaviour 139: 1029–1042.

14. Darling DC, Werren JH (1990) Biosystematics of *Nasonia* (Hymenoptera: Pteromalidae): two new species reared from birds’ nests in North America. Ann. Entomol. Soc. Amer. 83: 352–368.

15. Raychoudhury R, Desjardins CA, Buellesbach J, Loehlin DW, Grillenberger BK, et. al. (2010) Behavioral and genetic characteristics of a new species of *Nasonia*. Heredity 104: 278–288

16. Breeuwer JAJ, Werren JH (1990) Microorganisms associated with chromosome destruction and reproductive isolation between two insect species. Nature 346: 558–560.

17. Gadau J, Page Jr RE, Werren JH (1999) Mapping hybrid incompatibility loci in *Nasonia*. Genetics 153: 1731–1741.

18. Niehuis O, Judson AK, Gadau J (2008). Cytonuclear genic incompatibilities cause increased mortality in male F_2_ hybrids of *Nasonia giraulti* and *N vitripennis*. Genetics 178: 413–426.

19. Clark ME, O’Hara FP, Chawla A, Werren JH (2010). Behavioral and spermatogenic hybrid sterilty in *Nasonia*. Heredity 104: 289–301.

20. Koevoets T, Niehuis O, van den Zande L, Beukeboom LW (2012) Hybrid incompatibilities in the parasitic wasp genus *Nasonia*: negative effects of hemizygosity and the identification of transmission ratio distortion loci Heredity 108: 302–311.

21. van den Assem J (1986) Mating behaviour in parasitic wasps. pp. 137–167 in Insect Parasitoids, 13th Symposium of the Royal Entomological Society of London, edited by J. K. Waage, D. J. Greathead, Academic Press, London.

22. Rieseberg LH., Sinervo B, Linder CR, Ungerer MC, Arias M (1996) Role of gene interactions in hybrid speciation: evidence from ancient and experimental hybrids. Science 272: 741–745.

23. Pugh ARG, Ritchie MG (1996). Polygenic control of a mating signal in *Drosophila*. Heredity 77: 378–382.

24. Monforte A.J, Asins MJ, Carbonell EA (1997) Salt tolerance in *Lycopersicon* species. V. Does genetic variability at quantitative trait loci affect their analysis? Theor Appl Genet 95: 284–293.

25. True J R, Liu J, Stam LF, Zeng Z-B, Laurie CC (1997) Quantitative genetic analysis of divergence in male secondary sexual traits between *Drosophila simulans* and *Drosophila mauritiana*. Evolution 51: 816–832.

26. Hoikkala A, Päällysaho S, Aspi J, Lummer J (2000). Localization of genes affecting species differences in male courtship song between *Drosophila virilis* and *D*. *littoralis*. Genet Res Camb 75: 37–45.

27. Noor MAF, Grams KL, Bertucci LA, Almendarez Y, Reiland J, et al. (2001). The genetics of reproductive isolation and the potential for gene exchange between *Drosophila pseudoobscura* and *D*. *persimilis* via backcross hybrid males. Evolution 55: 512–521.

28. Gadau J, Page Jr RE, Werren JH (2002) The genetic basis of the interspecific differences in wing size in *Nasonia* (Hymenoptera; Pteromalidae): major quantitative trait loci and epistasis. Genetics 161: 673–684.

29. Moehring AJ, TFC. Mackay (2004) The quantitative genetic basis of male mating behavior in *Drosophila melanogaster*. Genetics 167: 1249–1263.

30. Moehring, AJ, Li J, Schug MD, Smith S, DeAngelis M et al. (2004) Quantitative trait loci for sexual isolation between *Drosophila simulans and D. mauritiana*. Genetics 167: 1265–1274.

31. Hoikkala H, Klappert K, Mazzi D (2005). Factors affecting male song evolution in *Drosophila montana*. Curr Top Dev Biol 67: 225–250.

32. Shaw KL, Parsons YM, Lesnick SC (2007). QTL analysis of a rapidly evolving speciation phenotype in the Hawaiian cricket *Laupala*. Mol Ecol 16: 2879–2892.

33. LH Rieseberg, MA Archer and RK Wayne 1999 Transgressive segregation, adaptation and speciation. Heredity 83:363–372.

34. Rieseberg LH, Raymond O, Roshenthal DM, Lai Z, Livingstone K et al. 2003 Major ecological transitions in wild sunflowers facilitated by hybridization. Science 301,1211– 1216.

35. Lexer C, Rosenthal DM, Raymond O, Donovan LA, Rieseberg, LH (2005) Genetics of species differences in the wild annual sunflowers, *Helianthus annus* and *H. petiolaris*. Genetics 169: 2225–2239.

36. Presgraves DC, Balagopalan L, Abmayr SM, Orr HA (2003) Adaptive evolution drives divergence of a hybrid inviability gene between two species of *Drosophila*. Nature 388: 663–666.

37. van den Assem J, Putters FA, van der Voort-Vinkestijn M-J (1984) Effects of exposure to an extremely low temperature on recovery of courtship behaviour after waning in the parasitic wasp *Nasonia vitripennis*. J Comp Physiol A 155: 233–237.

38. van den Assem J (1974) Male courtship patterns and female receptivity signal of Pteromalinae (Hym., Pteromalidae), with a consideration of some evolutionary trends and a comment on the taxonomic position of *Pachycrepoideus vindemiae*. Neth. J. Zool. 24: 253–278.

39. Gadau J, Heinze J, Hölldobler B, Schmid M (1996) Population and colony structure of the carpenter ant, *Camponotus floridanus*. Mol Ecol 5: 785–792.

40. Pietsch C, Rütten K, Gadau J (2004) Eleven microsatellite markers in *Nasonia*, Ashmead 1904 (Hymenoptera; Pteromalidae). Mol. Ecol. Notes 4: 43–45.

41. Beukeboom LW, Niehuis O, Pannebakker BA, Koevoets T, Gibson JD et al. (2010) A comparison of recombination frequencies in intraspecific versus interspecific mapping populations of *Nasonia*. Heredity. 104: 302–9.

42. Rütten KB, Pietsch C, Beukeboom L, Gadau J (2004) Chromsomal anchoring of behavioral and wing QTL in Nasonia. Cytogenet Genome Res 105: 126–133.

43. Werren JH, Richards S, Desjardins CA, Niehuis O, Gadau J et al. (2010). Functional and evolutionary insights from the genomes of three parasitoid *Nasonia* species. Science 327: 343–348.

44. Van Ooijen JW, 2004. MapQTL® 5, Software for the mapping of quantitative trait loci in experimental populations. Kyazma B.V., Wageningen, Netherlands.

45. Churchill GA, Doerge RW (1994) Empirical threshold values for quantitative trait mapping. Genetics 138: 963–971.

46. Broman, KW, H Wu, S Sen, and GA Churchill, 2003 R/qtl: QTL mapping in experimental crosses. Bioinformatics 19: 889–890.

47. Kankare M, Salminen T, Laiho A, Vesala L, Hoikkala A (2010). Changes in gene expression linked with adult reproductive diapause in a northern malt fly species: a candidate gene microarray study. BMC Ecol. 10: 3.

48. Munoz-Torres M., Reese JT, Childers CP, Bennett AK, Sundaram JP, et al. (2011) Hymenoptera Genome Database: integrated community resources for insect species of the order Hymenoptera. Nucleic Acids Res 39: D658–D662.

49. Li H, (2011) A quick method to calculate QTL confidence interval. Journal of Genetics 90: 355–360.

50. Falconer and McKay T (1996) Introduction to Quantitative Genetics. 4^th^ Edition Longman Group Ltd. Essex, England.

51. Ritchie MG, Phillips SDF (1998) The genetics of sexual isolation. In: Endless Forms: Species and Speciation (D. A. Howard & S. Berlocher, eds), pp. 291±308. Oxford University Press, Oxford.

52. Stern DL, Orgogozo V (2009) Is genetic evolution predictable? Science 323:746–751.

53. Orr HA (2005) The genetic theory of adaptation: a brief history. Nat Rev Genet 6: 119–127.

54. Marygold SJ, Leyland PC, Seal RL, Goodman JL, Thurmond JR, et al. (2013) FlyBase: improvements to the bibliography. Nucleic Acids Res 41(D1): D751–D757.

55. Niehuis O, Buellesbach J, Gibson JD, Pothmann D, Hanner C, et al. (2012) Behavioural and genetic analyses of *Nasonia* shed light on the evolution of sex pheromones. Nature 494: 345–348.

56. Gleason JM, Ritchie MG (2004). Do quantitative trait loci (QTL) for a courtship song difference between *Drosophila simulans* and *D*. *sechellia* coincide with candidate genes and intraspecific QTL? Genetics 166: 1303–1311.

57. Ruther J, Steiner S, Garbe LA (2008) 4-methylquinazoline is a minor component of the male sex pheromone in *Nasonia vitripennis*. J Chem Ecol. 34: 99–102.

58. Niehuis O, Büllesbach J, Judson AK, Schmitt T, Gadau J (2011) Genetics of cuticular hydrocarbon differences between males of the parasitoid wasps *Nasonia giraulti* and *Nasonia vitripennis*. Heredity 107: 61–70.

