## Supplementary material for "Genetic architecture of male courtship behavior differences in the parasitoid wasp genus *Nasonia* (Hymenoptera; Pteromalidae)": SOM Table1+2+3

**SOM Table 1:** Phenotypic Correlations (Spearman’s rank correlation coefficient) between all courtship components of our QTL analysis. All cycle and headnod phenotypes are highly correlated.

| | | | latency | fix-nod | cycle1 | cycle2 | cycle3 | cycle4 | hns1 | hns2 | hns3 | hns4 | h2_h1 | totalseries |
|---|---|---|---|---|---|---|---|---|---|---|---|---|---|---|
| Spearman's rho | latency | Correlation Coefficient | 1,000 | ,227$^{**}$ | ,217$^{**}$ | ,283$^{**}$ | ,295$^{**}$ | ,279$^{**}$ | -,028 | -,009 | ,056 | ,087 | ,031 | -,033 |
| | | Sig. (2-tailed) | . | ,000 | ,000 | ,000 | ,000 | ,000 | ,653 | ,882 | ,365 | ,168 | ,621 | ,640 |
| | | N | 268 | 236 | 255 | 257 | 210 | 199 | 266 | 267 | 266 | 254 | 265 | 202 |
| | fix-nod | Correlation Coefficient | | 1,000 | ,528$^{**}$ | ,525$^{**}$ | ,492$^{**}$ | ,462$^{**}$ | ,081 | ,086 | ,082 | ,083 | ,019 | -,117 |
| | | Sig. (2-tailed) | | . | ,000 | ,000 | ,000 | ,000 | ,185 | ,158 | ,179 | ,184 | ,752 | ,115 |
| | | N | | 270 | 257 | 258 | 193 | 182 | 269 | 269 | 269 | 255 | 268 | 184 |
| | cycle1 | Correlation Coefficient | | | 1,000 | ,830$^{**}$ | ,778$^{**}$ | ,702$^{**}$ | ,315$^{**}$ | ,305$^{**}$ | ,299$^{**}$ | ,246$^{**}$ | -,008 | -,133 |
| | | Sig. (2-tailed) | | | . | ,000 | ,000 | ,000 | ,000 | ,000 | ,000 | ,000 | ,891 | ,066 |
| | | N | | | 294 | 281 | 199 | 191 | 292 | 293 | 291 | 277 | 291 | 192 |
| | cycle2 | Correlation Coefficient | | | | 1,000 | ,833$^{**}$ | ,812$^{**}$ | ,293$^{**}$ | ,358$^{**}$ | ,372$^{**}$ | ,302$^{**}$ | ,092 | -,244$^{**}$ |
| | | Sig. (2-tailed) | | | | . | ,000 | ,000 | ,000 | ,000 | ,000 | ,000 | ,116 | ,001 |
| | | N | | | | 292 | 206 | 194 | 290 | 292 | 291 | 278 | 290 | 198 |
| | cycle3 | Correlation Coefficient | | | | | 1,000 | ,889$^{**}$ | ,221$^{**}$ | ,284$^{**}$ | ,381$^{**}$ | ,307$^{**}$ | ,040 | -,203$^{**}$ |
| | | Sig. (2-tailed) | | | | | . | ,000 | ,001 | ,000 | ,000 | ,000 | ,566 | ,004 |
| | | N | | | | | 210 | 195 | 210 | 209 | 210 | 201 | 209 | 195 |
| | cycle4 | Correlation Coefficient | | | | | | 1,000 | ,232$^{**}$ | ,269$^{**}$ | ,368$^{**}$ | ,341$^{**}$ | ,019 | -,213$^{**}$ |
| | | Sig. (2-tailed) | | | | | | . | ,001 | ,000 | ,000 | ,000 | ,787 | ,003 |
| | | N | | | | | | 199 | 199 | 198 | 199 | 199 | 198 | 186 |
| | Headnod 1 | Correlation Coefficient | | | | | | | 1,000 | ,720$^{**}$ | ,605$^{**}$ | ,592$^{**}$ | -,327$^{**}$ | -,113 |
| | | Sig. (2-tailed) | | | | | | | . | ,000 | ,000 | ,000 | ,000 | ,111 |
| | | N | | | | | | | 306 | 305 | 303 | 289 | 305 | 202 |
| | Headnod 2 | Correlation Coefficient | | | | | | | | 1,000 | ,754$^{**}$ | ,729$^{**}$ | ,357$^{**}$ | -,050 |
| | | Sig. (2-tailed) | | | | | | | | . | ,000 | ,000 | ,000 | ,480 |
| | | N | | | | | | | | 307 | 304 | 290 | 305 | 201 |
| | Headnod 3 | Correlation Coefficient | | | | | | | | | 1,000 | ,826$^{**}$ | ,208$^{**}$ | -,101 |
| | | Sig. (2-tailed) | | | | | | | | | . | ,000 | ,000 | ,154 |
| | | N | | | | | | | | | 305 | 290 | 302 | 202 |
| | Headnod 4 | Correlation Coefficient | | | | | | | | | | 1,000 | ,178$^{**}$ | -,078 |
| | | Sig. (2-tailed) | | | | | | | | | | . | ,002 | ,277 |
| | | N | | | | | | | | | | 291 | 288 | 194 |
| | h2_h1 | Correlation Coefficient | | | | | | | | | | | 1,000 | ,094 |
| | | Sig. (2-tailed) | | | | | | | | | | | . | ,185 |
| | | N | | | | | | | | | | | 305 | 201 |

| | | | | | | | | | | | | | | |
|---|---|---|---|---|---|---|---|---|---|---|---|---|---|---|
| | totalseries | Correlation Coefficient | | | | | | | | | | | | 1,000 |
| | | Sig. (2-tailed) | | | | | | | | | | | | . |
| | | N | | | | | | | | | | | | 202 |

**. Correlation is significant at the 0.01 level (2-tailed).

**SOM Table 2: Summary of the explained phenotypic variance from the two way interaction analysis.** For this analysis we conducted a whole genome two-way interaction analysis in which we searched for loci that had no significant effect on the phenotypes on their own ( see Fig. 3 secondary loci for a graphic depiction of all interactions) but had a significant phenotypic effect in connection with another locus. The column expl.var.add. represents the phenotypic variance explained by primary and secondary loci whose effect is additive. The column expl. var. epi is the phenotypic variance in our F2 hybrid mapping population that can be explained by non-additve (= epistatic) interactions between either primary or secondary QTL loci (Figure 3). The last column is the sum of the two previous columns and represents the phenotypic variance in our mapping population that can be explained by additive and epistatic interacting QTL.

| trait | expl. var. add. | expl. var. epi. | expl. var. total |
|---|---|---|---|
| latency | 4.97 | 0 | 4.97 |
| fix-nod | 32.06 | 2.20 | 34.26 |
| cycle1 | 39.61 | 8.71 | 48.31 |
| cycle2 | 33.86 | 2.65 | 36.51 |
| cycle3 | 27.56 | 1.44 | 29.00 |
| cycle4 | 18.94 | 0.00 | 18.94 |
| hnd1 | 21.63 | 0.00 | 21.63 |
| hnd2 | 29.43 | 0.00 | 29.43 |
| hnd3 | 29.95 | 0.00 | 29.95 |
| hnd4 | 23.63 | 0.00 | 23.63 |
| h2_h1 | 4.77 | 0.00 | 4.77 |
| minusnods | 12.29 | 3.16 | 15.45 |
| Forfeet rubbing | 6.93 | 1.85 | 8.78 |

**SOM Table 3:** Phenotypes of males from both parental species N. longicornis and N. vitripennis.

| species | fix1stnod | hnd1 | hnd2 | hnd3 | hnd4 | hndtot | cycel1 | cycle2 | cycle3 | h2_h1 |
|---|---|---|---|---|---|---|---|---|---|---|
| N. vitripennis | | 4 | | | 2 | | | | | |

| | | | | | | | | | | |
|---|---|---|---|---|---|---|---|---|---|---|
| N. vitripennis | | 7 | 5 | 5 | 5 | 7 | 7 | | | -2 |
| N. vitripennis | | 5 | 4 | 5 | 6 | 7 | | | | -1 |
| N. vitripennis | | 6 | 5 | 7 | 8 | 5 | | | | -1 |
| N. vitripennis | | 10 | 6 | 7 | 8 | 5 | 9 | | | -4 |
| N. vitripennis | | 6 | | | | | | | | |
| N. vitripennis | | 6 | 4 | | | | 8 | | | -2 |
| N. vitripennis | | 8 | | | | | 10 | | | |
| N. vitripennis | | 7 | 7 | | | | 10 | | | 0 |
| N. vitripennis | | 5 | 5 | 3 | 5 | 6 | 10 | | | 0 |
| N. vitripennis | | 6 | 3 | 4 | 5 | 7 | 8 | | | -3 |
| N. vitripennis | | 4 | 3 | 4 | 5 | 7 | 9 | | | -1 |
| N. vitripennis | | 6 | 4 | 5 | 5 | 7 | 6 | | | -2 |
| N. vitripennis | | 4 | 3 | 4 | 4 | 9 | 8 | | | -1 |
| N. vitripennis | | 4 | 3 | 4 | 5 | 10 | 7 | | | -1 |
| N. vitripennis | | 3 | 4 | 4 | 4 | 8 | 6 | | | 1 |
| N. vitripennis | | 4 | 3 | 4 | 5 | 7 | 7 | | | -1 |
| N. vitripennis | | 4 | 4 | | | | 6 | | | 0 |
| N. vitripennis | | 4 | 2 | 3 | 4 | 6 | 9 | | | -2 |
| N. vitripennis | | 4 | 4 | 3 | 4 | 13 | 4 | | | 0 |
| N. vitripennis | | 4 | 4 | 5 | 4 | 11 | 5 | | | 0 |
| N. vitripennis | | 4 | 4 | 4 | 6 | 9 | 6 | | | 0 |
| N. vitripennis | | 5 | | | | | 8 | | | |
| N. vitripennis | | 4 | 4 | | | | 6 | | | 0 |
| N. vitripennis | | 5 | 4 | | | | 8 | | | -1 |
| N. vitripennis | | 4 | 6 | 4 | | | 7 | | | 2 |
| N. vitripennis | | 4 | 2 | | | | 5 | | | -2 |
| N. vitripennis | | 5 | 4 | | | | 7 | | | -1 |

| | | | | | | | | | | |
|---|---|---|---|---|---|---|---|---|---|---|
| N. vitripennis | | 6 | 4 | | | | 8 | | | -2 |
| N. vitripennis | | 7 | 6 | | | | 10 | | | -1 |
| N. vitripennis | | 4 | 3 | 3 | 5 | 5 | 6 | | | -1 |
| N. vitripennis | | 5 | 4 | | | | 10 | | | -1 |
| N. vitripennis | | | | | | | | | | |
| N. vitripennis | | 6 | 4 | | | | 11 | | | -2 |
| N. vitripennis | | 6 | | | | | 11 | | | |
| N. vitripennis | | 6 | 4 | | | | 11 | | | -2 |
| N. vitripennis | | 4 | | | | | 7 | | | |
| N. vitripennis | | 5 | | | | | 7 | | | |
| N. vitripennis | | 6 | | | | | 7 | | | |
| N. vitripennis | | 5 | 3 | | | | 7 | | | -2 |
| N. vitripennis | | 5 | | | | | 7 | | | |
| N. vitripennis | | 5 | 4 | 4 | 4 | 8 | 8 | | | -1 |
| N. vitripennis | | 5 | 4 | 4 | 5 | 7 | 8 | | | -1 |
| N. vitripennis | | 4 | 3 | 3 | | | 6 | | | -1 |
| N. vitripennis | | 4 | 3 | 2 | 5 | 8 | 6 | | | -1 |
| N. vitripennis | | 4 | 4 | 4 | 5 | 7 | 8 | | | 0 |
| N. vitripennis | | 6 | 3 | 5 | 6 | 7 | 8 | | | -3 |
| N. vitripennis | | 4 | 4 | 5 | 4 | 6 | 5 | | | 0 |
| N. vitripennis | | 5 | 4 | 5 | 5 | 7 | 8 | | | -1 |
| N. vitripennis | | 5 | 4 | | | | 5 | | | -1 |
| N. vitripennis | | 4 | 4 | 4 | 4 | 6 | 8 | | | 0 |
| N. vitripennis | | 3 | 3 | 4 | 4 | 10 | 8 | | | 0 |
| N. vitripennis | | 5 | 4 | 5 | 4 | 8 | 7 | | | -1 |
| N. vitripennis | | 4 | 4 | 4 | 4 | 6 | 7 | | | 0 |
| N. vitripennis | | 4 | 4 | 4 | 4 | 8 | 7 | | | 0 |

| | | | | | | | | | | |
|---|---|---|---|---|---|---|---|---|---|---|
| N. vitripennis | | 5 | 3 | 4 | 4 | 7 | 6 | | | -2 |
| N. vitripennis | | 5 | 4 | 4 | 4 | 7 | 7 | | | -1 |
| N. vitripennis | | 3 | 2 | 3 | 3 | 7 | 6 | | | -1 |
| N. vitripennis | | 5 | 4 | 4 | 4 | 8 | 8 | | | -1 |
| N. vitripennis | | 5 | 4 | 4 | 5 | 8 | 5 | | | -1 |
| N. vitripennis | | 7 | 5 | 4 | 4 | 7 | 5 | | | -2 |
| N. vitripennis | | 7 | 5 | 6 | 5 | 6 | 5 | | | -2 |
| N. vitripennis | | 5 | 4 | 4 | 5 | 7 | | | | -1 |
| N. vitripennis | | 6 | 4 | 4 | 5 | 14 | 11 | | | -2 |
| N. vitripennis | | 8 | 4 | 4 | 5 | 6 | 8 | | | -4 |
| N. vitripennis | | 5 | 3 | 5 | 5 | 6 | 9 | | | -2 |
| N. vitripennis | | 3 | 4 | 4 | 5 | 8 | 6 | | | 1 |
| N. vitripennis | | 4 | 4 | 4 | 4 | 7 | 9 | | | 0 |
| N. vitripennis | | 7 | 4 | 7 | 8 | 7 | 7 | | | -3 |
| N. vitripennis | | 5 | 4 | 5 | 5 | 7 | 8 | | | -1 |
| N. vitripennis | | 6 | 5 | 5 | 6 | 8 | 8 | | | -1 |
| N. vitripennis | | 5 | 5 | 6 | 5 | 7 | 8 | | | 0 |
| N. vitripennis | | 5 | 3 | 5 | 5 | 6 | 9 | | | -2 |
| N. vitripennis | | 4 | 3 | 3 | 3 | 9 | | | | -1 |
| N. vitripennis | | | | | | | | | | |
| N. vitripennis | | 4 | 2 | 2 | 4 | 7 | 5 | | | -2 |
| N. vitripennis | | 3 | 2 | 3 | 3 | 5 | 4 | | | -1 |
| N. vitripennis | | 4 | 4 | 3 | 2 | 11 | | | | 0 |
| N. vitripennis | | 3 | 2 | 3 | 4 | 15 | 5 | | | -1 |
| N. vitripennis | | | | | | | | | | |
| N. vitripennis | 1 | 5 | 4 | 4 | 3 | | 6 | 6 | 7 | -1 |
| N. vitripennis | 1 | 7 | 7 | 8 | 8 | | 8 | 10 | 10 | 0 |

| | | | | | | | | | | |
|---|---|---|---|---|---|---|---|---|---|---|
| N. vitripennis | | 7 | 7 | | | | 11 | | | 0 |
| N. vitripennis | 2 | 5 | 4 | 6 | | | 6 | 6 | | -1 |
| N. vitripennis | 2 | 4 | | | | | 5 | | | |
| N. vitripennis | 1 | 5 | 4 | 5 | 5 | | 5 | 5 | 6 | -1 |
| N. vitripennis | 2 | 5 | | | | | 8 | | | |
| N. vitripennis | 1 | 6 | 5 | 6 | 7 | | 8 | 8 | 8 | -1 |
| N. vitripennis | 6 | 5 | 6 | 6 | 7 | | 7 | 8 | 8 | 1 |
| N. vitripennis | 2 | 7 | 6 | 6 | 7 | | 7 | 7 | 8 | -1 |
| N. vitripennis | 1 | 5 | 5 | | | | 8 | 8 | | 0 |
| N. vitripennis | 1 | 5 | 5 | 7 | 7 | | 5 | 7 | 7 | 0 |
| N. vitripennis | 2 | 6 | 6 | 7 | 6 | | 8 | 7 | 10 | 0 |
| N. vitripennis | 2 | 7 | 6 | 5 | 6 | | 8 | 8 | 11 | -1 |
| N. vitripennis | 1 | 5 | 6 | 7 | 4 | | 5 | 7 | 8 | 1 |
| N. vitripennis | | 5 | 3 | 3 | 4 | | 4 | 4 | 6 | -2 |
| N. vitripennis | 1 | 5 | 3 | | | | 5 | 6 | | -2 |
| N. vitripennis | | | | | | | | | | |
| N. vitripennis | 12 | 3 | 4 | 5 | 5 | | 6 | 7 | 7 | 1 |
| N. vitripennis | 2 | 5 | 4 | 4 | 6 | | 6 | 6 | 8 | -1 |
| N. vitripennis | 1 | 5 | 3 | 4 | 5 | | 5 | 5 | 5 | -2 |
| N. vitripennis | 5 | 6 | 4 | 5 | 5 | | 6 | 5 | 7 | -2 |
| N. vitripennis | 2 | 8 | 5 | 6 | 5 | | 8 | 7 | 9 | -3 |
| N. vitripennis | | | | | | | | | | |
| N. vitripennis | 1 | 3 | 4 | 5 | 7 | | | | 9 | 1 |
| N. vitripennis | 1 | 5 | 4 | 4 | 3 | | 5 | 5 | 6 | -1 |
| N. vitripennis | 2 | 7 | 5 | 5 | | | 7 | 7 | 7 | -2 |
| N. vitripennis | 1 | 3 | 4 | 4 | 6 | | 4 | 5 | 6 | 1 |
| N. vitripennis | 6 | 6 | 5 | 5 | 5 | | 7 | 7 | 8 | -1 |

| | | | | | | | | | | |
|---|---|---|---|---|---|---|---|---|---|---|
| N. vitripennis | | 7 | 6 | 6 | 7 | | | | | -1 |
| N. vitripennis | 1 | 6 | 7 | 5 | 6 | | 7 | 8 | 8 | 1 |
| N. vitripennis | 1 | 6 | 4 | 4 | 4 | | 6 | 8 | 9 | -2 |
| N. vitripennis | 1 | 3 | 6 | 8 | 7 | | 8 | 7 | 7 | 3 |
| N. vitripennis | 8 | 6 | | | | | 6 | | | |
| N. vitripennis | 2 | 7 | 6 | 6 | 7 | | 9 | 7 | 8 | -1 |
| N. vitripennis | | | 5 | 8 | 7 | | | 6 | 9 | |
| N. vitripennis | 1 | 6 | 4 | 5 | 5 | | 6 | 7 | 7 | -2 |
| N. vitripennis | 5 | 3 | 3 | 5 | 5 | | 6 | 6 | 7 | 0 |
| N. vitripennis | 1 | 2 | 4 | | | | 9 | 8 | | 2 |
| N. vitripennis | 2 | 5 | 5 | 4 | 6 | | 6 | 5 | 6 | 0 |
| N. vitripennis | | | 3 | 3 | 6 | | | 6 | 7 | |
| N. vitripennis | | | 6 | 7 | 6 | | 6 | 7 | 8 | |
| N. vitripennis | 1 | 5 | 6 | 5 | 5 | | 7 | 7 | 8 | 1 |
| N. vitripennis | 1 | 5 | | | | | 5 | | | |
| N. vitripennis | | 5 | 3 | | 5 | | 7 | 6 | 7 | -2 |
| N. vitripennis | 1 | 4 | 4 | 4 | 1 | | 5 | 6 | 6 | 0 |
| N. vitripennis | | 6 | 5 | 5 | 5 | | 6 | 7 | 7 | -1 |
| N. vitripennis | 1 | 5 | 3 | 4 | 4 | | 7 | 6 | 7 | -2 |
| N. vitripennis | 6 | 5 | 6 | 5 | 5 | | 6 | 7 | 8 | 1 |
| N. vitripennis | 1 | | 5 | 4 | | | 6 | 7 | 8 | |
| N. vitripennis | 1 | 6 | 4 | 5 | | | 5 | 5 | 7 | -2 |
| N. vitripennis | 2 | 3 | 4 | 5 | 4 | | 5 | 6 | 6 | 1 |
| N. vitripennis | | | | | | | | | | |
| N. vitripennis | | | 5 | 5 | 6 | | | 7 | 8 | |
| N. vitripennis | 9 | 5 | 5 | 4 | 4 | | 8 | 7 | 9 | 0 |
| N. vitripennis | | | | | | | | | | |

| | | | | | | | | | | |
|---|---|---|---|---|---|---|---|---|---|---|
| N. vitripennis | 1 | 4 | 3 | 5 | 5 | | 5 | 5 | 7 | -1 |
| N. vitripennis | 1 | 5 | | | | | 6 | | | |
| N. vitripennis | 1 | 5 | | | | | 7 | | | |
| N. vitripennis | | 5 | 5 | 5 | 6 | | | | | |
| N. longicornis | | 6 | 4 | | | | 19 | | | -2 |
| N. longicornis | | 2 | 3 | 2 | 3 | 10 | 12 | | | 1 |
| N. longicornis | | 2 | 2 | 4 | 4 | 9 | 11 | | | 0 |
| N. longicornis | | 1 | 1 | 3 | 3 | 12 | 12 | | | 0 |
| N. longicornis | | 2 | 4 | 5 | | | 17 | | | 2 |
| N. longicornis | | 2 | 3 | 4 | 4 | 11 | 13 | | | 1 |
| N. longicornis | | 1 | 2 | 2 | 3 | 12 | 11 | | | 1 |
| N. longicornis | | 1 | 1 | 2 | 3 | 11 | 11 | | | 0 |
| N. longicornis | | 7 | | | | | 21 | | | |
| N. longicornis | | 1 | 2 | 3 | 3 | 9 | 10 | | | 1 |
| N. longicornis | | 1 | 2 | 2 | 4 | 17 | 9 | | | 1 |
| N. longicornis | | 1 | 2 | 2 | 2 | 20 | 11 | | | 1 |
| N. longicornis | | | | | | | | | | |
| N. longicornis | | 2 | 2 | 2 | 3 | 9 | 13 | | | 0 |
| N. longicornis | | 1 | 2 | 2 | 3 | 12 | 14 | | | 1 |
| N. longicornis | | 3 | | | | | 12 | | | |
| N. longicornis | | 1 | 2 | 3 | 3 | 5 | 14 | | | 1 |
| N. longicornis | | 2 | 2 | 3 | 3 | 14 | 12 | | | 0 |
| N. longicornis | | 1 | 3 | 4 | | 4 | 14 | | | 2 |
| N. longicornis | | 2 | 1 | | | | 11 | | | -1 |
| N. longicornis | | 2 | 2 | 2 | 2 | 8 | 11 | | | 0 |
| N. longicornis | | 1 | 2 | | | | 13 | | | 1 |
| N. longicornis | | 1 | 2 | 2 | 4 | 9 | 12 | | | 1 |

| | | | | | | | | | | |
|---|---|---|---|---|---|---|---|---|---|---|
| N. longicornis | | 2 | 2 | 2 | 2 | 6 | 14 | | | 0 |
| N. longicornis | | 2 | 3 | 4 | 5 | 8 | 19 | | | 1 |
| N. longicornis | | 2 | 2 | 2 | 2 | 5 | 13 | | | 0 |
| N. longicornis | | 1 | 2 | 2 | 3 | 11 | 10 | | | 1 |
| N. longicornis | | 1 | 2 | | | | 12 | | | 1 |
| N. longicornis | | 1 | 2 | 2 | 2 | 7 | 12 | | | 1 |
| N. longicornis | | 1 | 1 | 2 | 2 | 8 | 12 | | | 0 |
| N. longicornis | | 3 | 4 | 4 | 4 | 6 | 12 | | | 1 |
| N. longicornis | | 1 | 2 | 2 | 4 | 7 | 9 | | | 1 |
| N. longicornis | | 2 | 2 | 2 | 2 | 6 | 13 | | | 0 |
| N. longicornis | | 2 | 2 | 2 | 3 | 9 | 15 | | | 0 |
| N. longicornis | | 1 | 1 | 2 | 2 | 8 | 13 | | | 0 |
| N. longicornis | | 2 | 2 | 3 | 4 | 7 | 17 | | | 0 |
| N. longicornis | | | | | | | | | | |
| N. longicornis | | 1 | 1 | 2 | 2 | 10 | 10 | | | 0 |
| N. longicornis | | 1 | 2 | 2 | 2 | 12 | 14 | | | 1 |
| N. longicornis | | 1 | 2 | | | | 11 | | | 1 |
| N. longicornis | | 2 | 2 | 3 | | | 16 | | | 0 |
| N. longicornis | | 1 | 2 | 3 | 3 | 6 | 11 | | | 1 |
| N. longicornis | | 1 | 2 | 2 | 2 | 7 | 13 | | | 1 |
| N. longicornis | | 1 | 1 | | | | 10 | | | 0 |
| N. longicornis | | 2 | 2 | 2 | 3 | 8 | 15 | | | 0 |
| N. longicornis | | 1 | 1 | | | | 13 | | | 0 |
| N. longicornis | | 1 | 1 | 2 | 2 | 5 | 13 | | | 0 |
| N. longicornis | | 1 | 1 | 2 | | | 15 | | | 0 |
| N. longicornis | | 1 | 1 | 2 | 2 | 9 | 13 | | | 0 |
| N. longicornis | | 1 | | | | | 22 | | | |

| | | | | | | | | | | |
|---|---|---|---|---|---|---|---|---|---|---|
| N. longicornis | | 1 | 1 | 2 | 2 | 8 | 14 | | | 0 |
| N. longicornis | | 1 | 1 | 2 | 2 | 17 | 12 | | | 0 |
| N. longicornis | | 2 | 2 | 3 | 3 | 9 | 15 | | | 0 |
| N. longicornis | | 1 | 1 | 3 | 2 | 8 | 12 | | | 0 |
| N. longicornis | | 1 | 1 | 1 | 1 | 5 | 12 | | | 0 |
| N. longicornis | | 1 | 1 | 3 | 2 | 8 | 12 | | | 0 |
| N. longicornis | | | | | | | | | | |
| N. longicornis | | | | | | | | | | |
| N. longicornis | | 1 | 1 | 2 | 2 | 9 | 13 | | | 0 |
| N. longicornis | | 1 | 3 | 4 | 5 | 9 | | | | 2 |
| N. longicornis | | 3 | 3 | 3 | 4 | 7 | 17 | | | 0 |
| N. longicornis | | 1 | 1 | 2 | 3 | 7 | 15 | | | 0 |
| N. longicornis | | 1 | 2 | 2 | 3 | 8 | 18 | | | 1 |
| N. longicornis | | 3 | 3 | 6 | 7 | 6 | 14 | | | 0 |
| N. longicornis | | 1 | 1 | 2 | 2 | 12 | 11 | | | 0 |
| N. longicornis | | 1 | 2 | 2 | 2 | 14 | 14 | | | 1 |
| N. longicornis | | 1 | 1 | 1 | 2 | 11 | 11 | | | 0 |
| N. longicornis | | 1 | 1 | 2 | 2 | 14 | 13 | | | 0 |
| N. longicornis | | 1 | 1 | 1 | 2 | 10 | 12 | | | 0 |
| N. longicornis | | 2 | 3 | 4 | | | 10 | | | 1 |
| N. longicornis | | 2 | 2 | 2 | 2 | 6 | 12 | | | 0 |
| N. longicornis | | 1 | 1 | 3 | 5 | 6 | 10 | | | 0 |
| N. longicornis | | 2 | 3 | 3 | 3 | 8 | 12 | | | 1 |
| N. longicornis | | 1 | 1 | 2 | 3 | 6 | 12 | | | 0 |
| N. longicornis | | 1 | 1 | | | | 10 | | | 0 |
| N. longicornis | | 1 | 1 | | | | 10 | | | 0 |
| N. longicornis | | 2 | 2 | 2 | 3 | 7 | 11 | | | 0 |

| | | | | | | | | | | |
|---|---|---|---|---|---|---|---|---|---|---|
| N. longicornis | | 1 | 1 | 2 | 2 | 5 | 10 | | | 0 |
| N. longicornis | | 2 | 2 | 2 | 2 | 5 | 18 | | | 0 |
| N. longicornis | | 1 | 2 | 3 | 3 | 8 | 12 | | | 1 |
| N. longicornis | | 1 | 2 | 2 | 2 | 6 | 13 | | | 1 |
| N. longicornis | | 1 | 1 | 2 | 2 | 5 | 13 | | | 0 |
| N. longicornis | | 2 | 2 | 3 | 4 | 10 | 13 | | | 0 |
| N. longicornis | | 1 | 1 | 1 | 2 | 7 | 14 | | | 0 |
| N. longicornis | | 1 | 2 | 3 | 4 | 6 | 11 | | | 1 |
| N. longicornis | 6 | 1 | 2 | 2 | 2 | | 10 | 9 | 10 | 1 |
| N. longicornis | 6 | 3 | 2 | 3 | 4 | | 11 | 10 | 11 | -1 |
| N. longicornis | 7 | 1 | 2 | 3 | 3 | | 8 | 10 | 10 | 1 |
| N. longicornis | 17 | 1 | 1 | 2 | 2 | | 13 | 11 | 12 | 0 |
| N. longicornis | 16 | 1 | 1 | 2 | 3 | | 10 | 10 | 11 | 0 |
| N. longicornis | 9 | 2 | 1 | 2 | 2 | | 10 | 21 | 11 | -1 |
| N. longicornis | 9 | 1 | 2 | 2 | 2 | | 11 | 12 | 12 | 1 |
| N. longicornis | 6 | 2 | 1 | 2 | 1 | | 10 | 10 | 11 | -1 |
| N. longicornis | | | 1 | 2 | 3 | | | 10 | 10 | |
| N. longicornis | 7 | 1 | 2 | 2 | 3 | | 7 | 11 | 12 | 1 |
| N. longicornis | | | | | | | | | | |
| N. longicornis | 14 | 2 | 2 | 3 | 3 | | 13 | 13 | 14 | 0 |
| N. longicornis | 4 | 1 | 2 | 3 | 2 | | 11 | 12 | 12 | 1 |
| N. longicornis | | | | | | | | | | |
| N. longicornis | 8 | 2 | 3 | 4 | 3 | | 13 | 13 | 13 | 1 |
| N. longicornis | | | | | | | | | | |
| N. longicornis | 12 | 2 | 2 | 3 | 4 | | 14 | 13 | 15 | 0 |
| N. longicornis | | | | | | | | | | |
| N. longicornis | 52 | 4 | 2 | 4 | 5 | | 16 | 14 | 15 | -2 |

| | | | | | | | | | | |
|---|---|---|---|---|---|---|---|---|---|---|
| N. longicornis | 7 | 1 | 2 | 2 | 3 | | 11 | 13 | 14 | 1 |
| N. longicornis | | | | | | | | | | |
| N. longicornis | | | | | | | | | | |
| N. longicornis | 7 | 1 | 1 | 2 | 4 | | 12 | 11 | 11 | 0 |
| N. longicornis | 6 | 3 | 4 | 5 | 4 | | 11 | 12 | 13 | 1 |
| N. longicornis | 11 | 2 | 2 | 3 | 3 | | 11 | 10 | 12 | 0 |
| N. longicornis | | 1 | 2 | 4 | 3 | | | 11 | 14 | 1 |
| N. longicornis | 7 | 1 | 1 | 1 | 1 | | 9 | 9 | 10 | 0 |
| N. longicornis | 19 | 2 | 2 | | | | 13 | 13 | | 0 |
| N. longicornis | | | | | | | | | | |
| N. longicornis | 9 | 2 | 2 | 2 | 3 | | 10 | 11 | 11 | 0 |
| N. longicornis | | | | | | | | | | |
| N. longicornis | 4 | 1 | 2 | 2 | 3 | | 11 | 11 | 13 | 1 |
| N. longicornis | 5 | 1 | 2 | 2 | 2 | | 10 | 13 | 13 | 1 |
| N. longicornis | 10 | 1 | 2 | 3 | 3 | | 10 | 12 | 13 | 1 |
| N. longicornis | 6 | 2 | 2 | 2 | | | 14 | 12 | | 0 |
| N. longicornis | 5 | 1 | 2 | 2 | 1 | | 13 | 14 | 13 | 1 |
| N. longicornis | 11 | 2 | 2 | 2 | 3 | | 12 | 12 | 14 | 0 |
| N. longicornis | 8 | 1 | 2 | 3 | 4 | | 10 | 12 | 12 | 1 |
| N. longicornis | 6 | 1 | 2 | | | | 13 | 11 | | 1 |
| N. longicornis | | | | | | | | | | |
| N. longicornis | 5 | 1 | 2 | 2 | 3 | | 9 | 12 | 11 | 1 |
| N. longicornis | 9 | 1 | 1 | 1 | | | 12 | 11 | 12 | 0 |
| N. longicornis | 6 | 1 | 2 | 2 | 2 | | 12 | 11 | 11 | 1 |
| N. longicornis | 9 | 1 | 1 | 2 | 2 | | 10 | 10 | 11 | 0 |
| N. longicornis | 4 | 1 | 2 | 2 | 2 | | 8 | 11 | 12 | 1 |
| N. longicornis | 6 | 1 | 2 | 3 | 2 | | 10 | 12 | 12 | 1 |

| | | | | | | | | | | |
|---|---|---|---|---|---|---|---|---|---|---|
| N. longicornis | 4 | 1 | 1 | 4 | 1 | | 12 | 11 | 15 | 0 |
| N. longicornis | | | | | | | | | | |
| N. longicornis | 9 | 2 | 1 | 1 | 2 | | 15 | 12 | 12 | -1 |
| N. longicornis | 6 | 2 | 2 | 2 | 3 | | 12 | 12 | 13 | 0 |
| N. longicornis | 4 | 1 | 1 | 2 | 2 | | 8 | 10 | 11 | 0 |
| N. longicornis | 4 | 1 | 2 | 2 | 3 | | 10 | 12 | 12 | 1 |
| N. longicornis | 4 | 2 | 2 | 3 | 2 | | 10 | 12 | 11 | 0 |
| N. longicornis | 4 | 1 | 1 | 1 | 2 | | 10 | 10 | 11 | 0 |
| N. longicornis | 8 | 2 | 1 | | | | 14 | | | -1 |
| N. longicornis | 6 | 1 | 1 | 1 | 2 | | 13 | 12 | 12 | 0 |
| Average N. vitripennis | 2.3 | 5.1 | 4.2 | 4.7 | 5.0 | 7.6 | 7.0 | 6.6 | 7.6 | -0.8 |
| STDEV | 2.43 | 1.29 | 1.14 | 1.25 | 1.32 | 2.08 | 1.66 | 1.14 | 1.23 | 1.20 |
| Variance | 6.05 | 1.67 | 1.31 | 1.57 | 1.76 | 4.39 | 2.77 | 1.32 | 1.54 | 1.45 |
| Average N. longicornis | 8.7 | 1.5 | 1.8 | 2.4 | 2.7 | 8.7 | 12.3 | 11.6 | 12.1 | 0.3 |
| StDev | 7.48 | 0.89 | 0.72 | 0.90 | 1.00 | 3.17 | 2.52 | 1.85 | 1.36 | 0.69 |
| Variance | 57.25 | 0.80 | 0.52 | 0.82 | 1.00 | 10.19 | 6.38 | 3.51 | 1.89 | 0.48 |
