## Supplementary material for "Genetic architecture of male courtship behavior differences in the parasitoid wasp genus *Nasonia* (Hymenoptera; Pteromalidae)": SOM Figure 1

SOM Figure 1: **Effect of sample size on estimation of explained phenotypic variance (Beavis effect) for cycle time 1-4, headnod 1-4, latency and fix-nod.** For all traits the explained phenotypic variance (left panel) and LOD scores (right panel) is shown as a function of sample size based on a bootstrap analysis (1000 bootstraps/sample size). Boxplots show means/SD/min-max. In a significant number of permutations, QTL would go undetected because they fall below the genome wide detection limit of 1% if the number of individuals in a mapping population is lower than 200 (for most traits the 0.01 significance threshold is around LOD 2.7, dashed line in the second panel). As predicted, the variance in the estimation of a QTL effect becomes smaller as sample size goes up (left panel). A significant QTL in our studies should always be detected if the mapping population is larger than 250 individuals (all LOD scores are above the detection threshold). For each trait the location of the QTL under consideration is given as chromosome and position, i.e. Cycle 1 / 1_50.3 means that the analysis was focused on the cycle 1 QTL at position 50.3 (see table 2).

### Cycle 1 / I_50.3

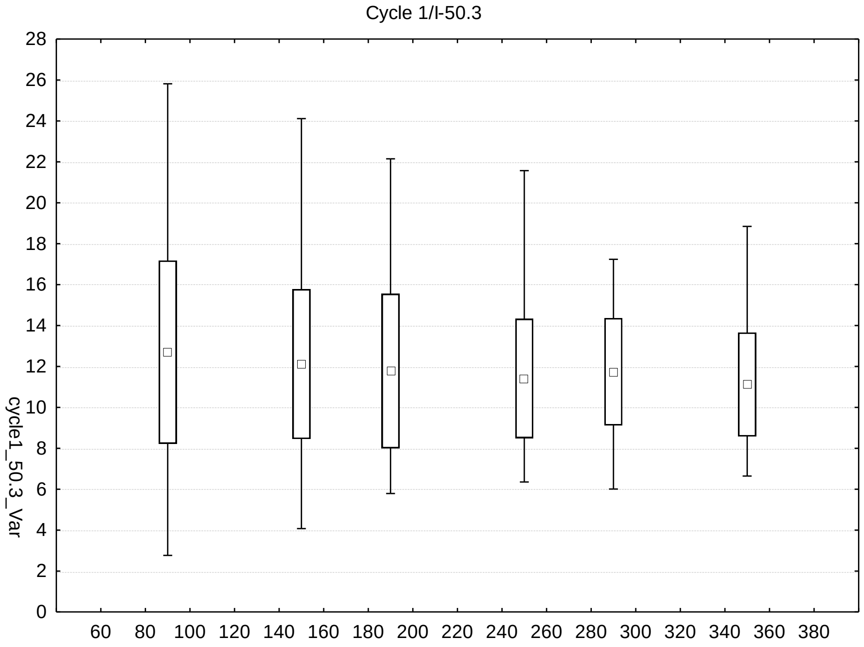


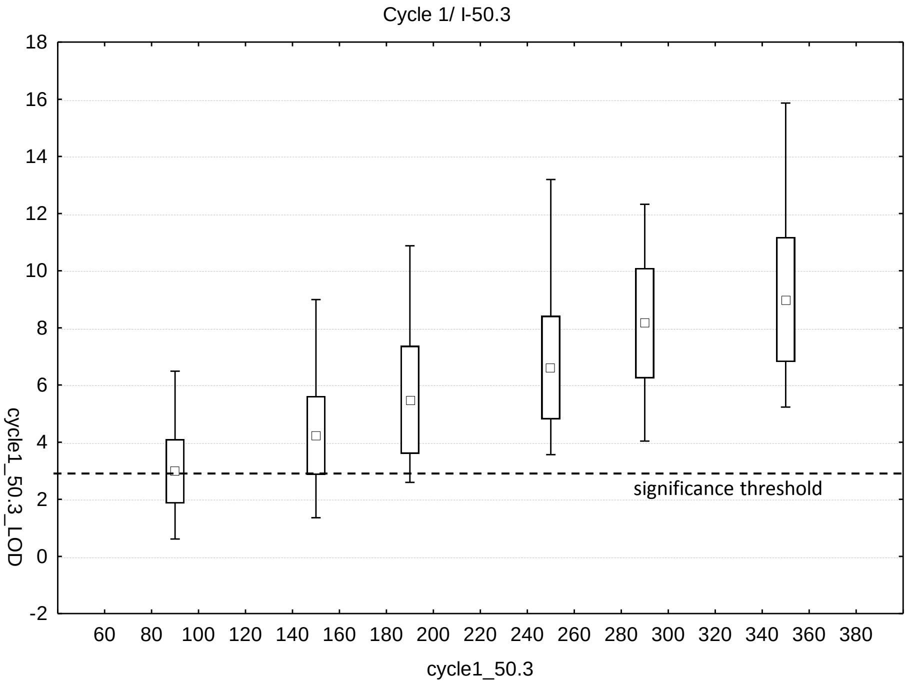


*Mean, Mean±SD (Box); Min-Max (Whiskers)

### Cycle 2 1_50.3

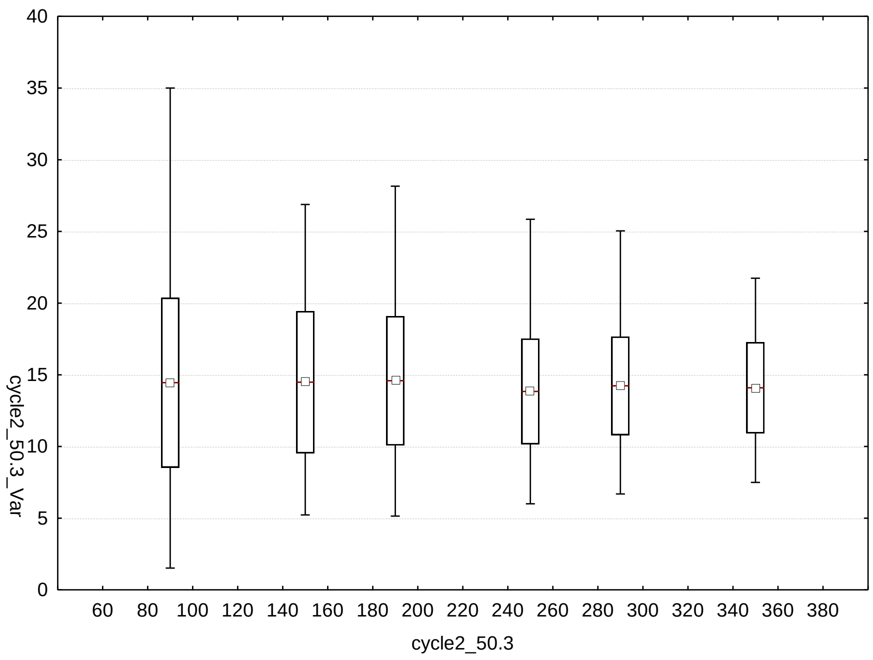


Variance explained

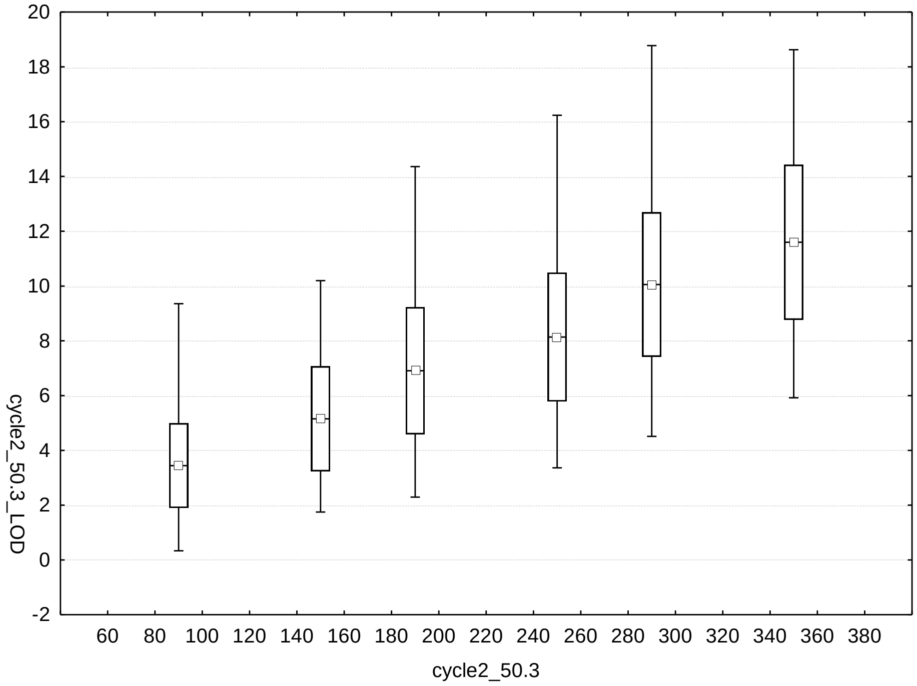


LOD scores

### Cycle 3 1_50.3

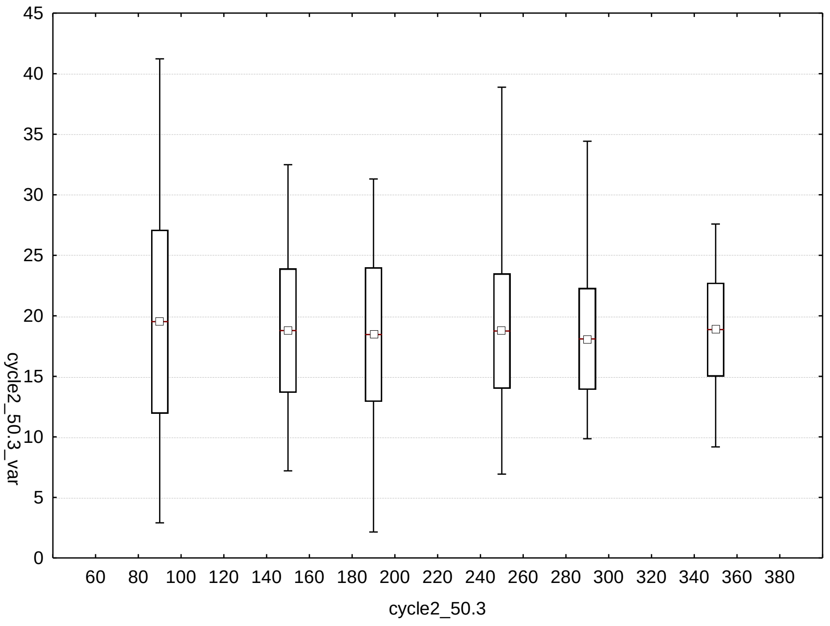

45
40
35
30
25
20
15
10
5
0
60 80 100 120 140 160 180 200 220 240 260 280 300 320 340 360 380
cycle2_50.3
cycle2_50.3_var


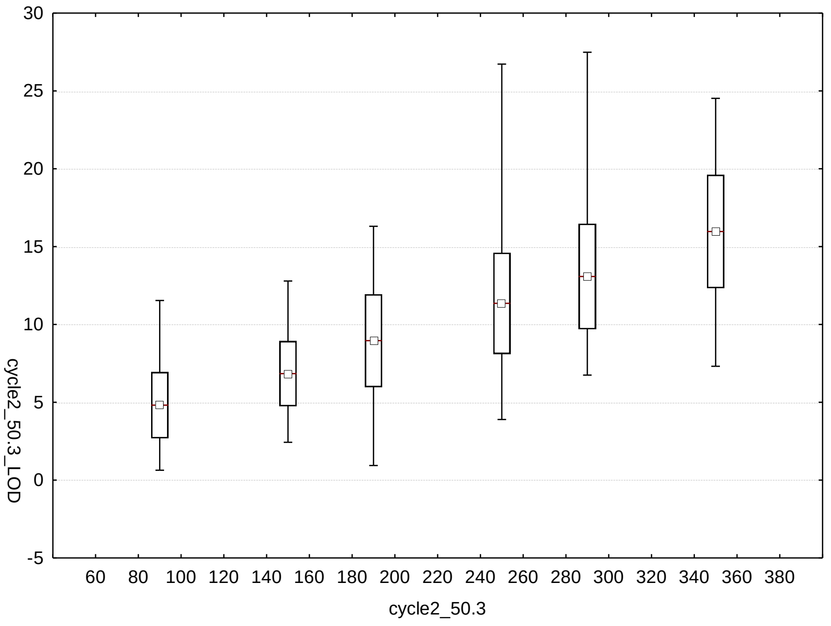

30
25
20
15
10
5
0
-5
60 80 100 120 140 160 180 200 220 240 260 280 300 320 340 360 380
cycle2_50.3
cycle2_50.3_LOD

### Cycle 4 1_50.3

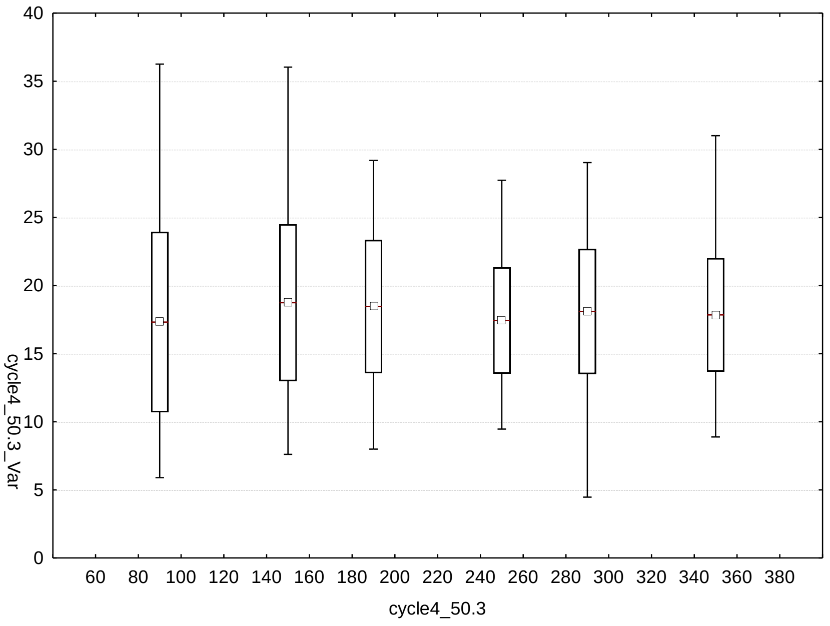

0
5
10
15
20
25
30
35
40
cycle4_50.3_Var
60 80 100 120 140 160 180 200 220 240 260 280 300 320 340 360 380
cycle4_50.3


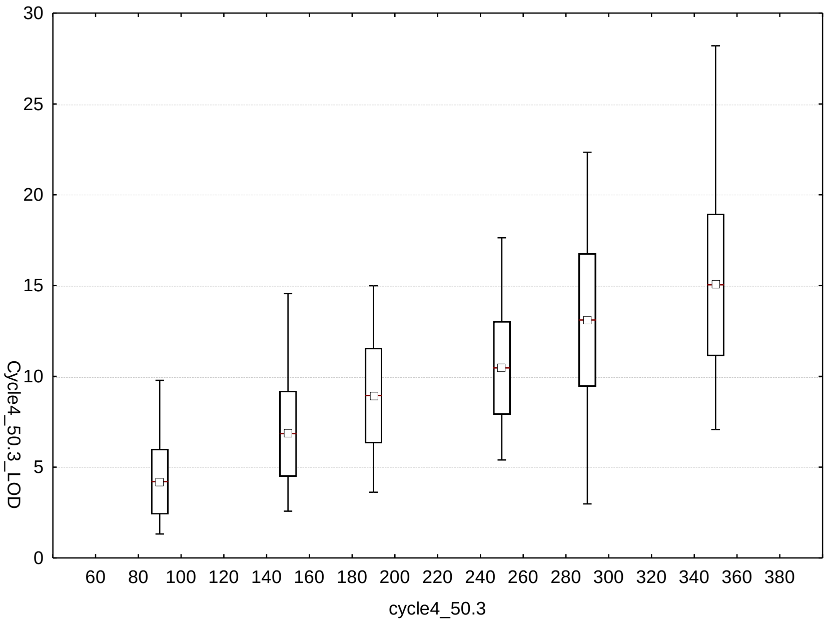

0
5
10
15
20
25
30
Cycle4_50.3_LOD
60 80 100 120 140 160 180 200 220 240 260 280 300 320 340 360 380
cycle4_50.3

### Headnod 1-III-66.3

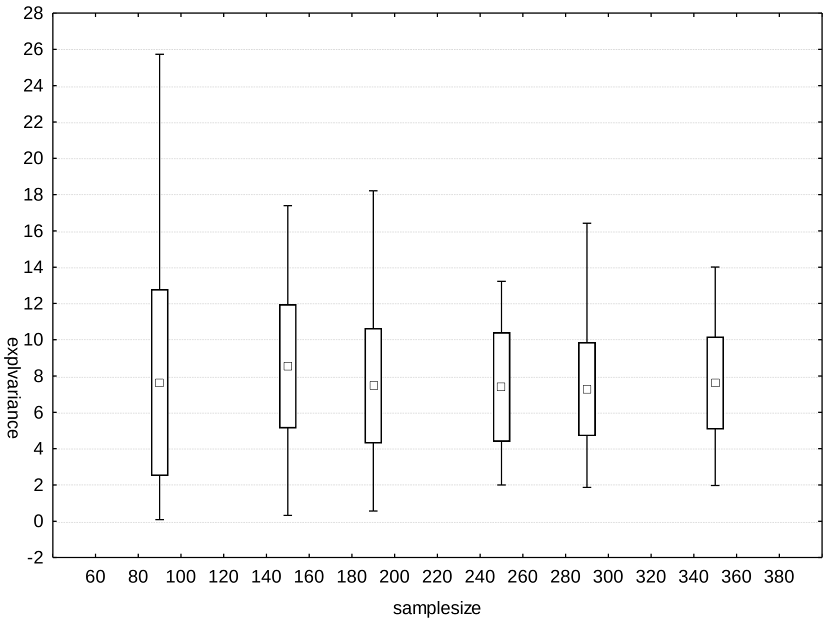


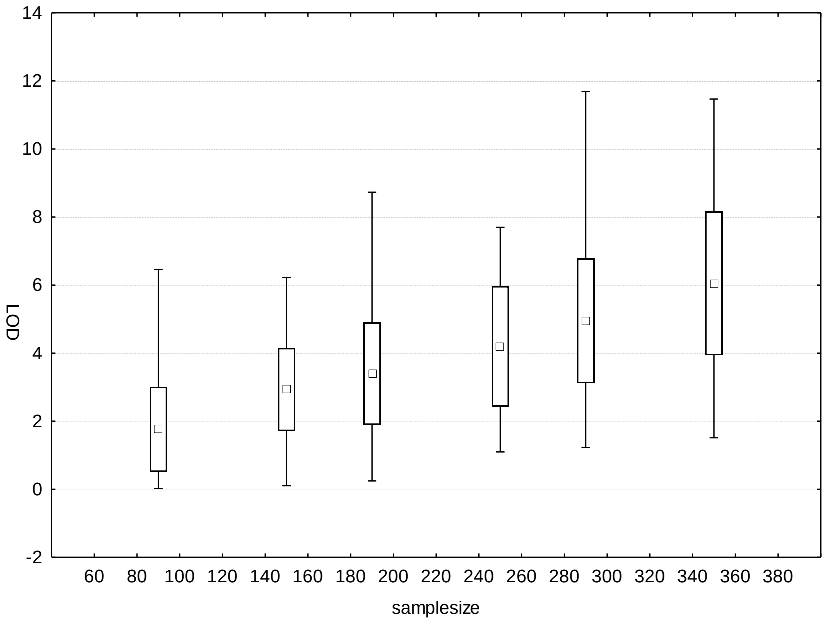

### Headnod 2 - III-66.3

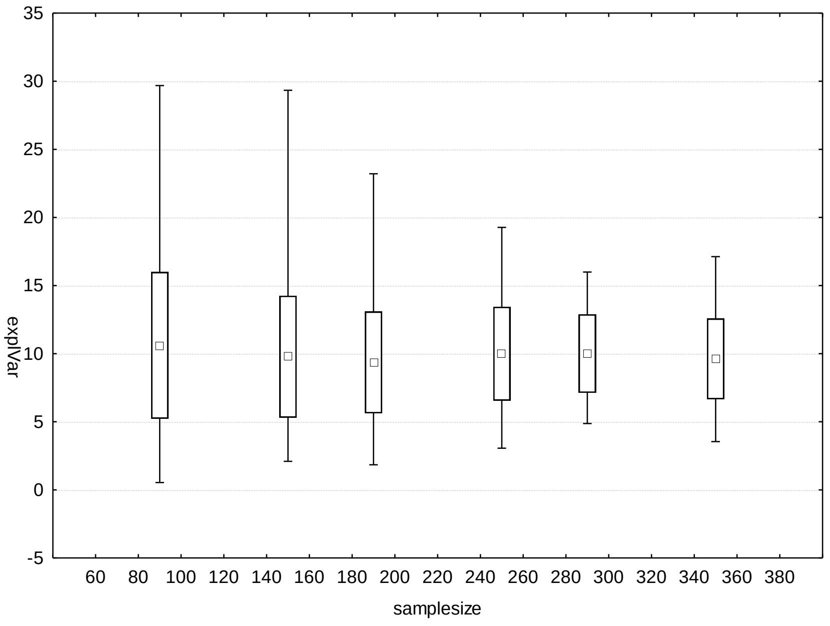


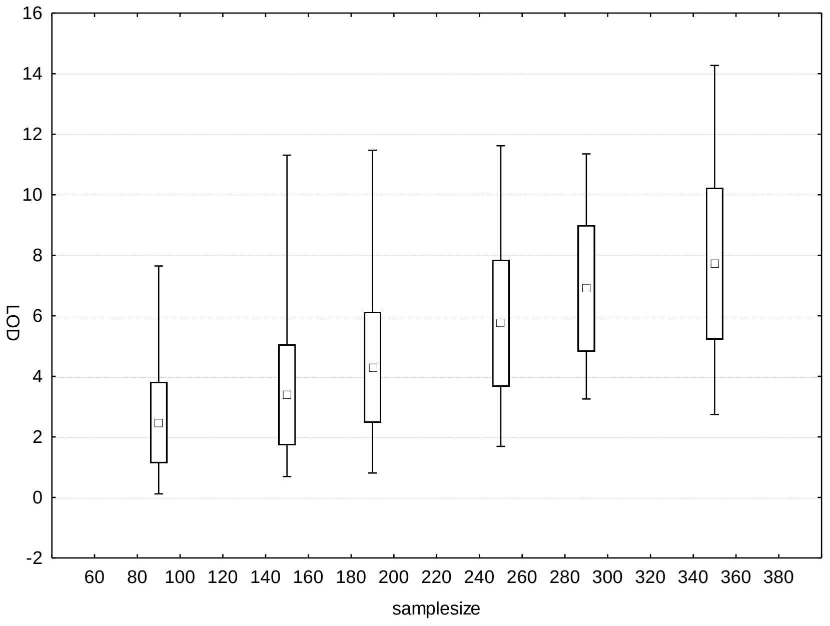

### Headnod 3 – III-66.3

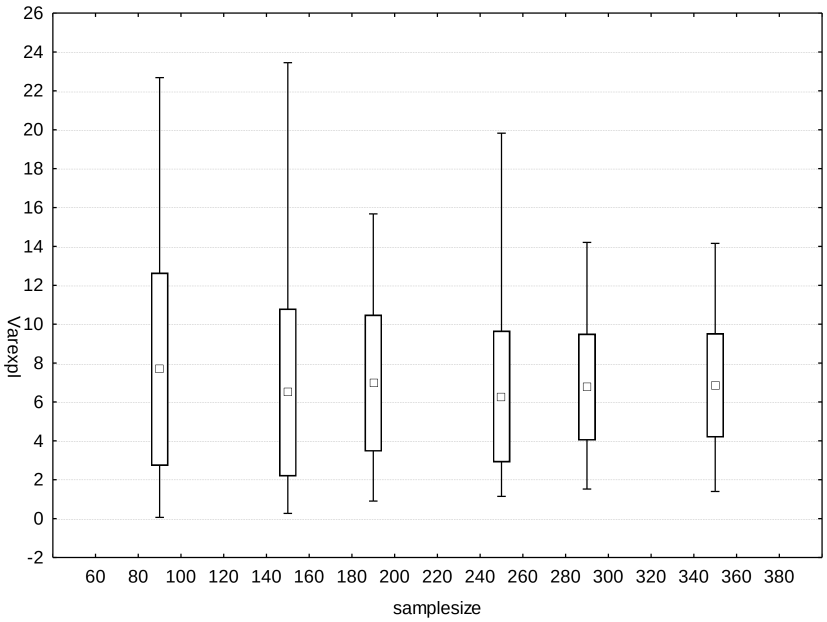


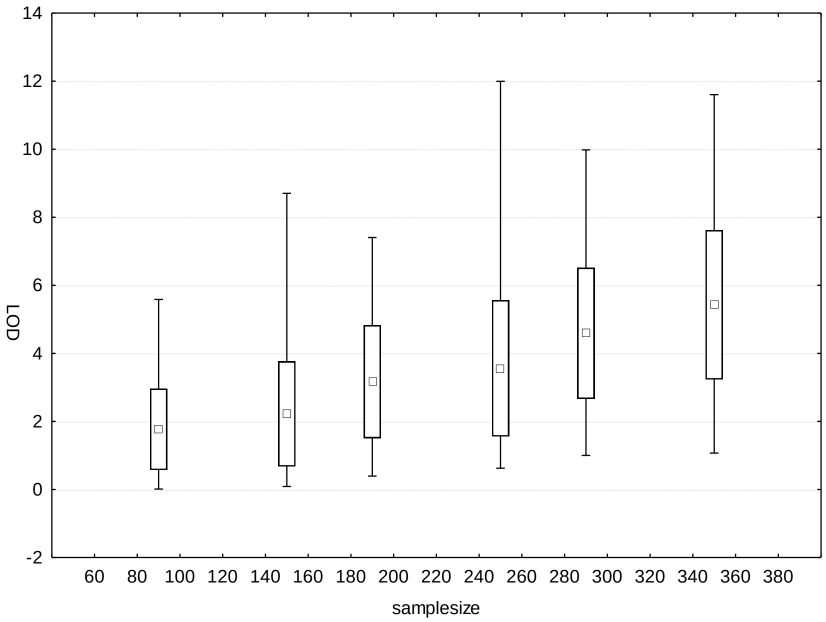

### Headnod 4 – III-66.3

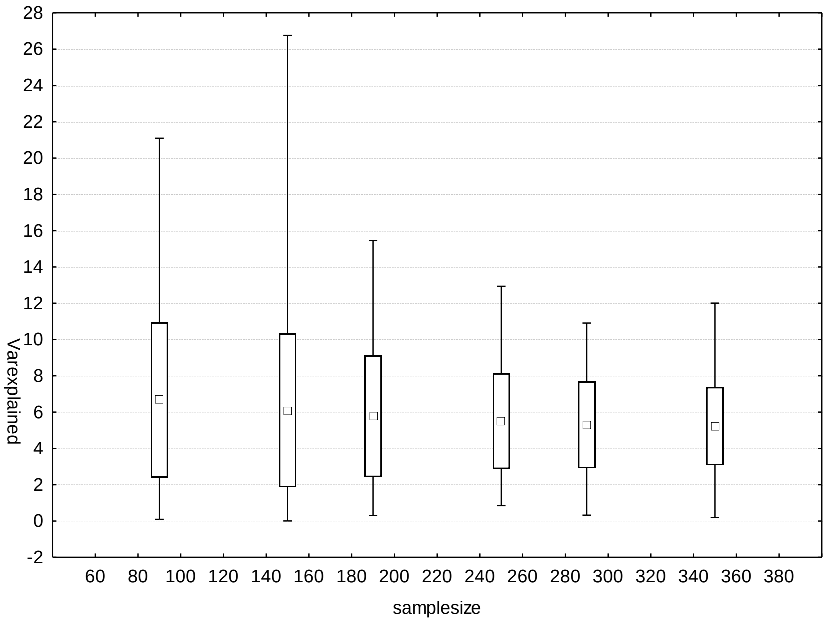


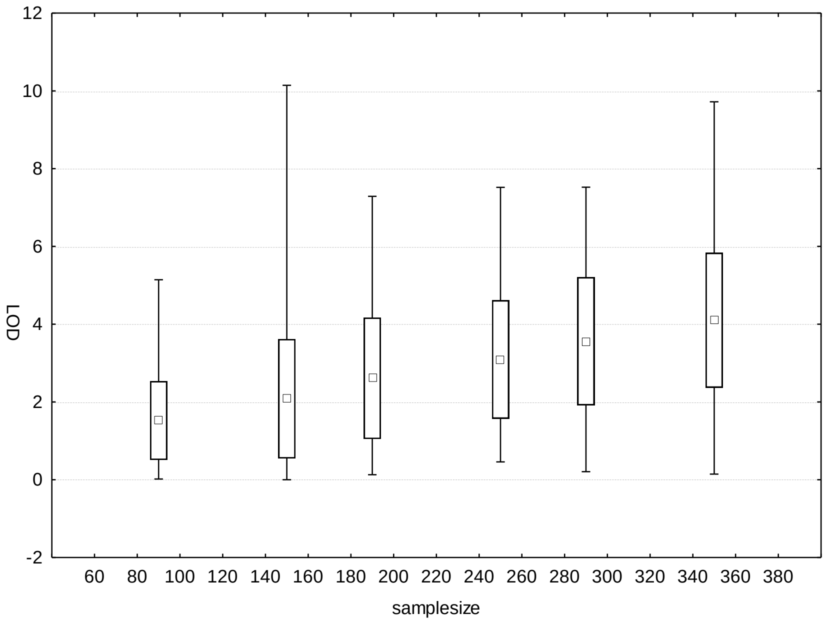

### Cycle 1 – V-55

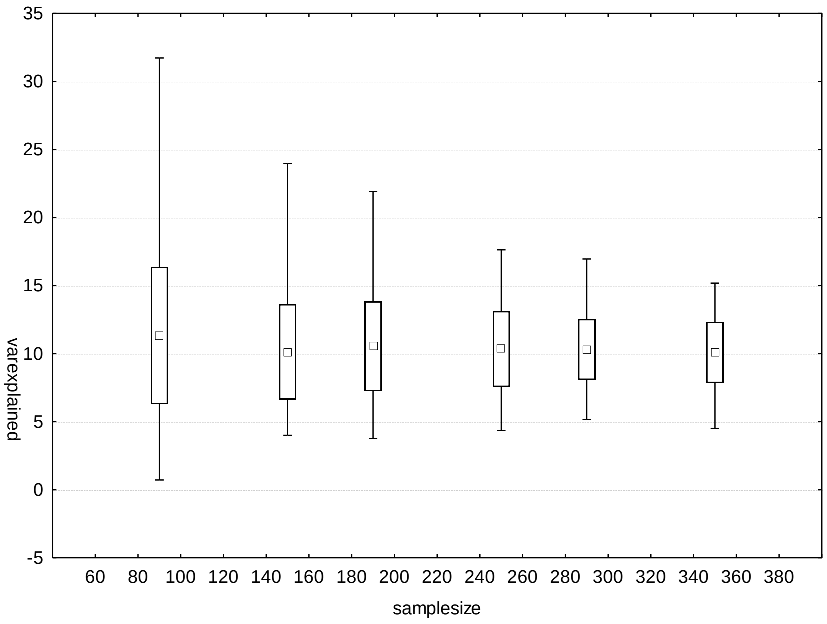

varexplained
samplesize
35
30
25
20
15
10
5
0
-5
60 80 100 120 140 160 180 200 220 240 260 280 300 320 340 360 380


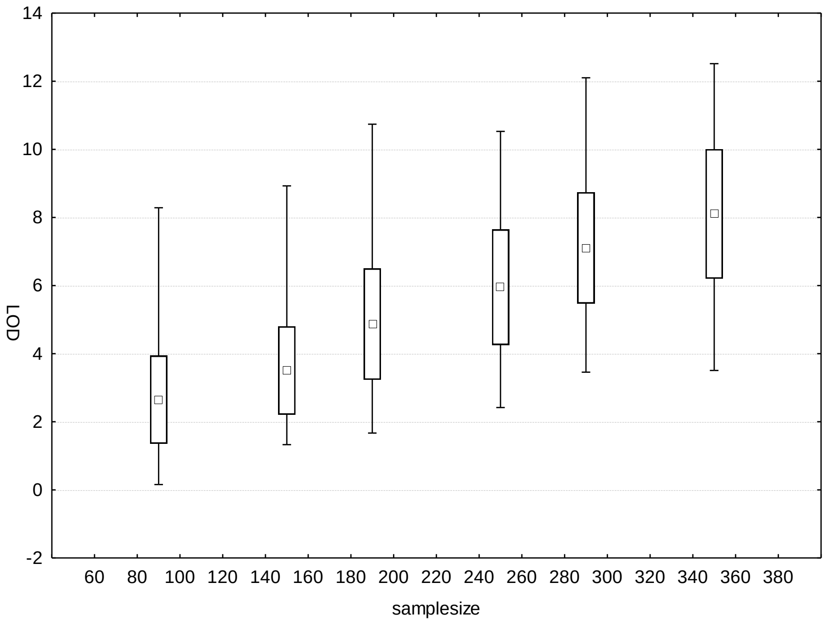

LOD
samplesize
14
12
10
8
6
4
2
0
-2
60 80 100 120 140 160 180 200 220 240 260 280 300 320 340 360 380

### Cycle 2 – V-55

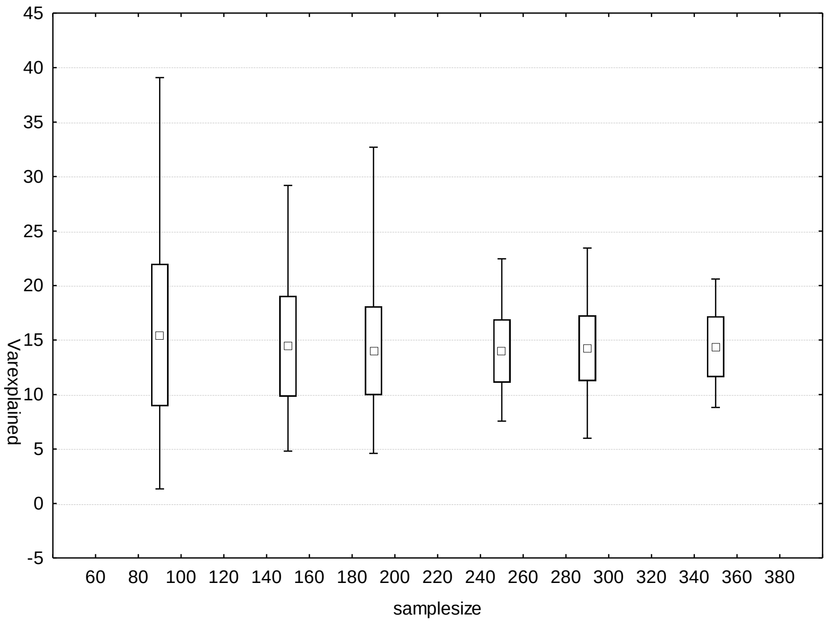

45
40
35
30
25
20
15
10
5
0
-5
Varexplained
60 80 100 120 140 160 180 200 220 240 260 280 300 320 340 360 380
samplesize


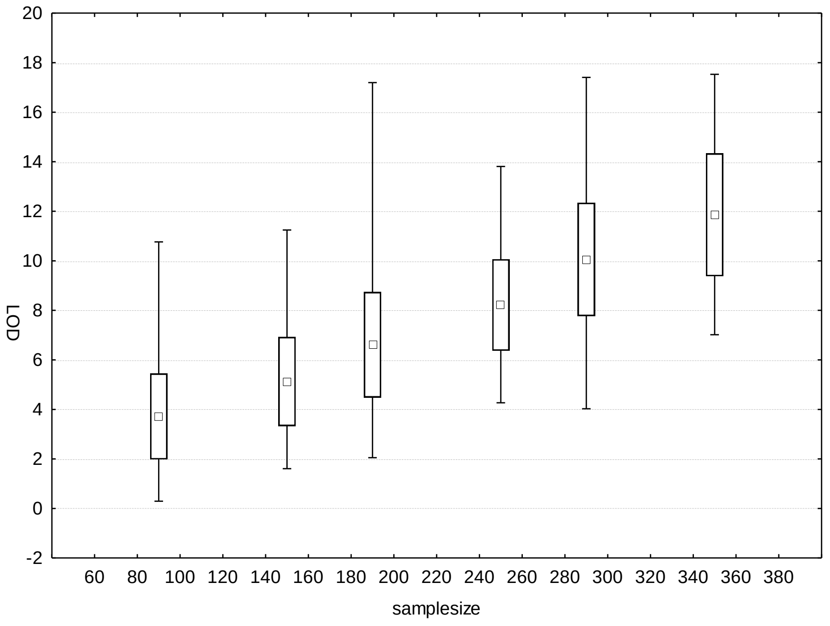

20
18
16
14
12
10
8
6
4
2
0
-2
LOD
60 80 100 120 140 160 180 200 220 240 260 280 300 320 340 360 380
samplesize

### Cycle 3 – V-55

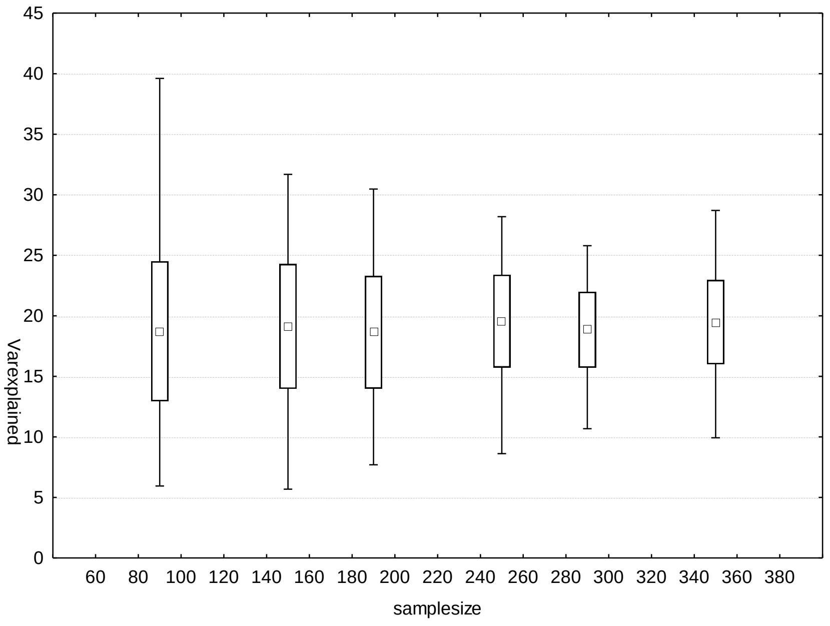
Varexplained
0
5
10
15
20
25
30
35
40
45
60 80 100 120 140 160 180 200 220 240 260 280 300 320 340 360 380
samplesize

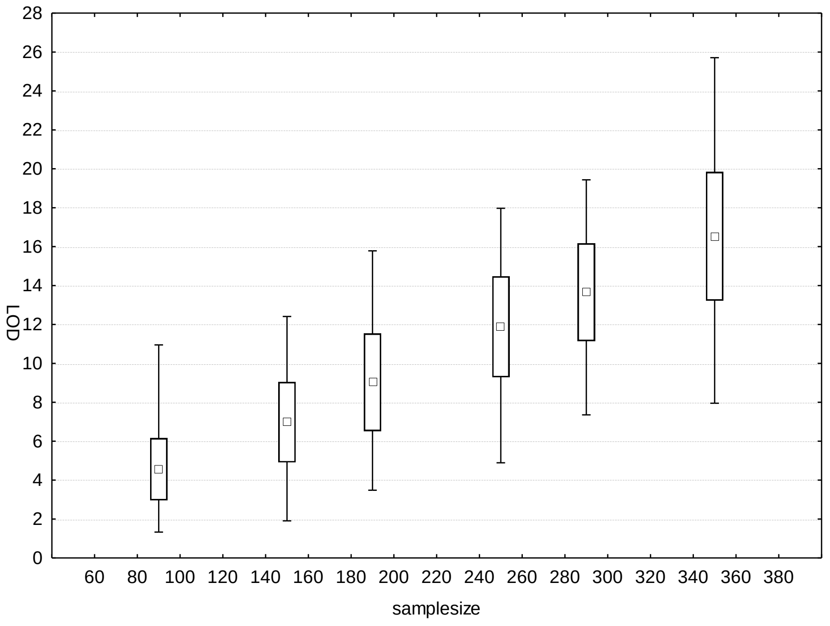
LOD
0
2
4
6
8
10
12
14
16
18
20
22
24
26
28
60 80 100 120 140 160 180 200 220 240 260 280 300 320 340 360 380
samplesize

### Cycle 4 – V-55

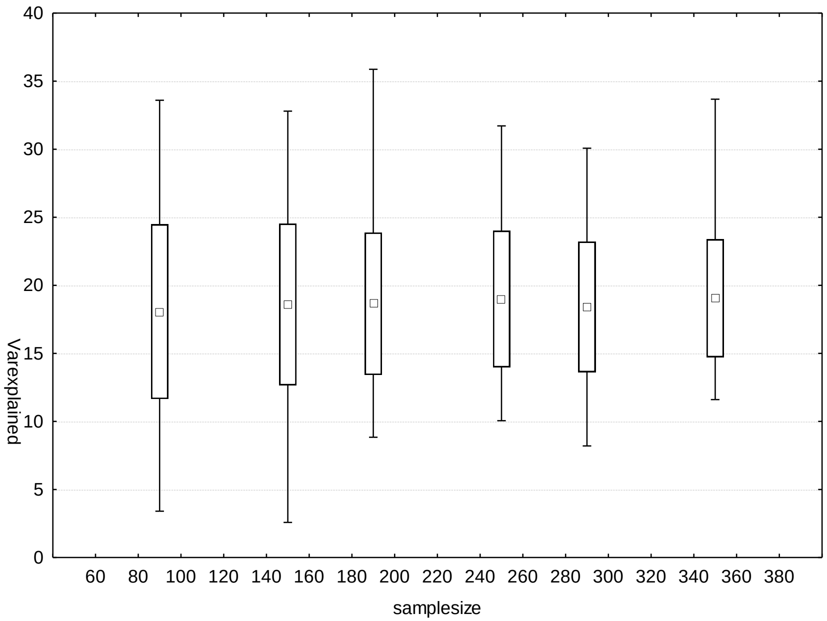

40
35
30
25
20
15
10
5
0
Varexplained
60 80 100 120 140 160 180 200 220 240 260 280 300 320 340 360 380
samplesize


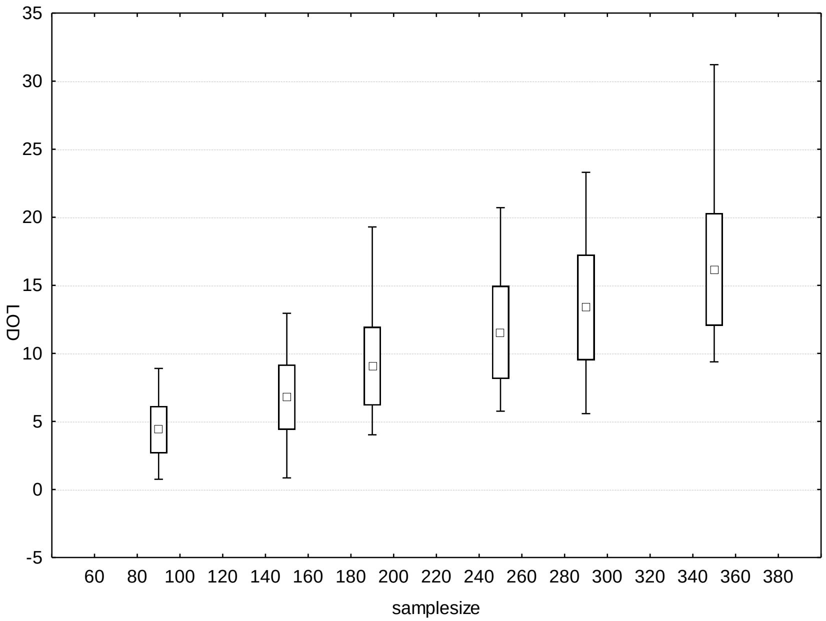

35
30
25
20
15
10
5
0
-5
LOD
60 80 100 120 140 160 180 200 220 240 260 280 300 320 340 360 380
samplesize

### Latency 1_50.3

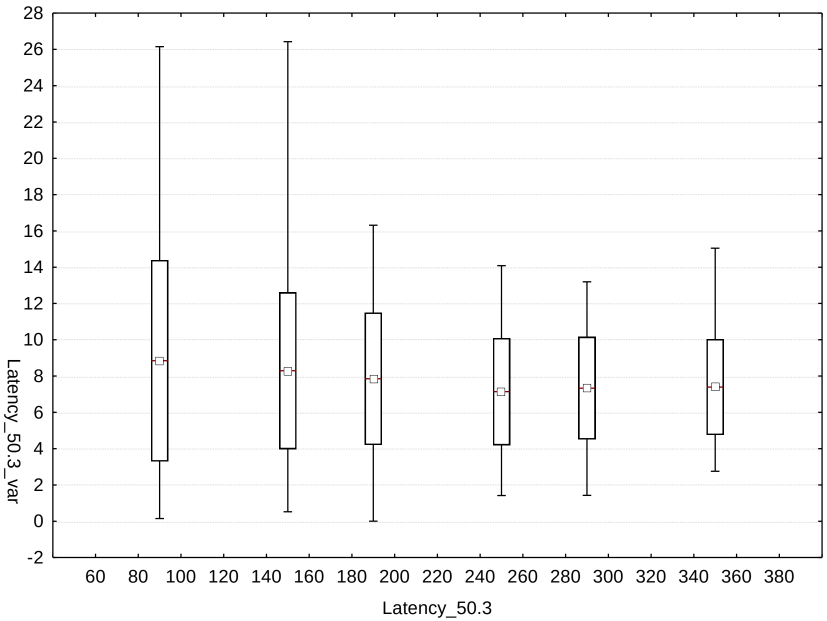


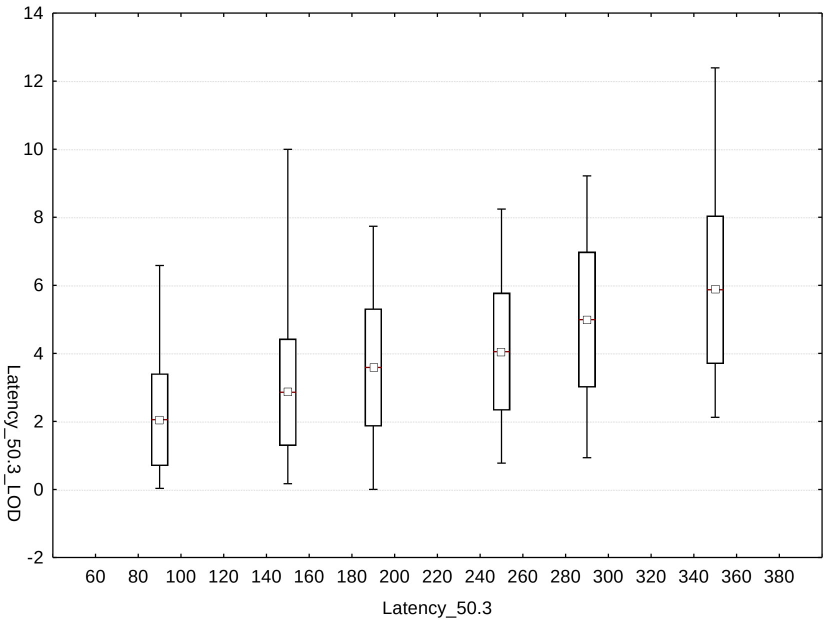

### Fix-Nod 1-37.1

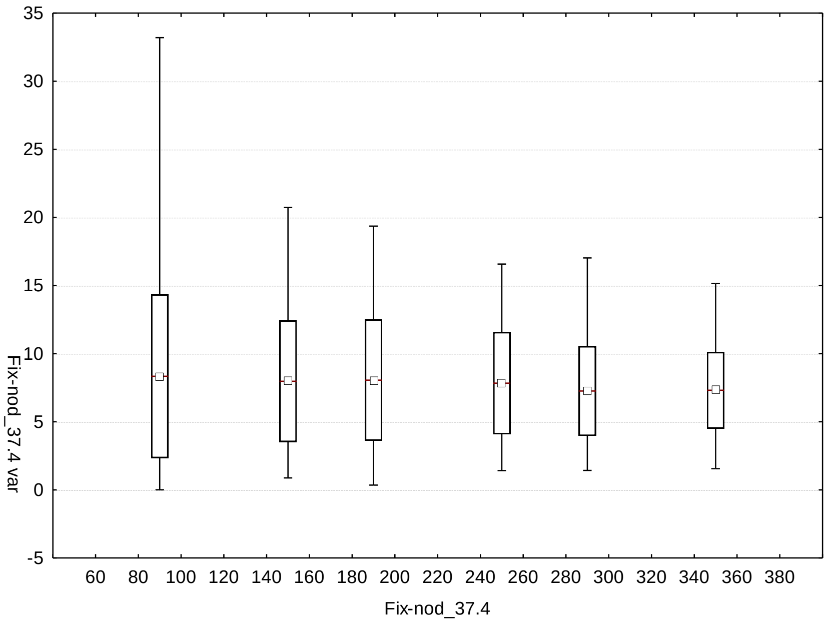

Fix-nod_37.4 var
35
30
25
20
15
10
5
0
-5
60 80 100 120 140 160 180 200 220 240 260 280 300 320 340 360 380
Fix-nod_37.4


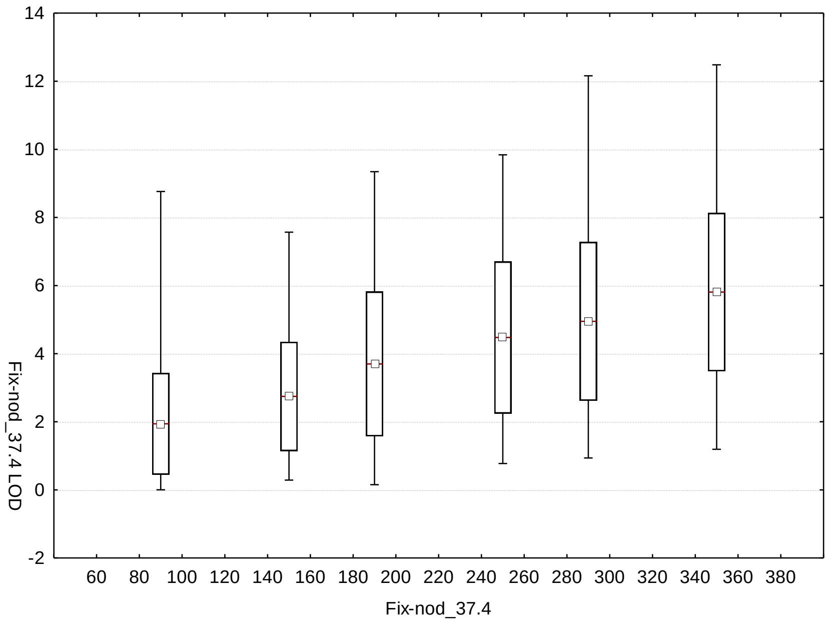

Fix-nod_37.4 LOD
14
12
10
8
6
4
2
0
-2
60 80 100 120 140 160 180 200 220 240 260 280 300 320 340 360 380
Fix-nod_37.4
